# The cellular and molecular landscape of hypothalamic patterning and differentiation

**DOI:** 10.1101/657148

**Authors:** Dong Won Kim, Parris Whitney Washington, Zoe Qianyi Wang, Sonia Hao Lin, Changyu Sun, Basma Taleb Ismail, Hong Wang, Lizhi Jiang, Seth Blackshaw

## Abstract

The hypothalamus is a central regulator of many innate behaviors essential for survival, but the molecular mechanisms controlling hypothalamic patterning and cell fate specification are poorly understood. To identify genes that control hypothalamic development, we have used single-cell RNA sequencing (scRNA-Seq) to profile mouse hypothalamic gene expression across 12 developmental time points between embryonic day 10 and postnatal day 45. This identified genes that delineated clear developmental trajectories for all major hypothalamic cell types, and readily distinguished major regional subdivisions of the developing hypothalamus. By using our developmental dataset, we were able to rapidly annotate previously unidentified clusters from existing scRNA-Seq datasets collected during development, and to identify the developmental origins of major neuronal populations of the ventromedial hypothalamus. We further show that our approach can rapidly and comprehensively characterize mutants that have altered hypothalamic patterning, identifying *Nkx2.1* as a negative regulator of prethalamic identity. These data serve as a resource for further studies of hypothalamic development, physiology and dysfunction.

## Introduction

The hypothalamus is comprised of a diverse array of neuronal and glial cell types, many of which are organized into spatially discrete clusters or nuclei^1–3^. Stereotactic lesion and focal stimulation studies have identified individual nuclei as essential for regulating a broad range of homeostatic physiological processes, ranging from circadian rhythms to hunger; behaviors such as mating, aggression and care of young; and cognitive processes such as motivation, reward, and memory^4–7^. More recently, opto- and chemogenetic techniques have made it possible to identify the role of individual hypothalamic neuronal subtypes in controlling some of these behaviors^8–10^.

Progress in this area has been hampered, however, by the fact that hypothalamic cell types thus far have remained quite poorly characterized, despite recent efforts aimed at using scRNA-Seq to classify cells in certain hypothalamic regions^11–14^.Still less is known about how hypothalamic cell types acquire their identities during development. Even the basic spatial organization of the developing hypothalamus, and its relationship to other forebrain structures such as the prethalamus and telencephalon, remains contentious^15–17^. Previous efforts using microarray analysis coupled with large-scale two-color *in situ* hybridization have identified a set of molecular markers that uniquely define spatial domains of the early embryonic hypothalamus and adjacent diencephalic regions^2^, while parallel efforts using high-throughput *in situ* hybridization have identified additional region-specific markers^18,19^.

These datasets have been used as the basis for genetic studies that selectively disrupt development of specific hypothalamic regions and/or cell types^20–24^, leading to identification of novel functions for previously characterized hypothalamic regions or cell types^25,26^. However, these datasets have important limitations: they do not provide cellular resolution of gene expression data, and they do not efficiently measure coexpression of multiple genes. In addition, despite the availability of many highly specific molecular markers, analysis of mutants that affect hypothalamic development is currently both slow and difficult, owing to the complexity of this structure.

Recent advances in single-cell RNA-Seq technology (scRNA-Seq)^27^ have made it possible to both analyze the development of complex organs at cellular resolution and to also rapidly and comprehensively characterize the molecular phenotype of developmental mutants^28^. In this study, we use scRNA-Seq to profile changes in gene expression and cell composition across the full course of mouse hypothalamic development, with a particular focus on identifying genes that control glial differentiation and function. We next focus on identifying genes that control hypothalamic regionalization and neurogenesis in the early embryo, and integrate these findings to generate a *Hy*pothalamic *D*evelopmental *D*atabase (HyDD), which identifies selective markers of each region of the developing hypothalamus and prethalamus. We next use the HyDD to rapidly annotate cell types in previously published scRNA-Seq datasets, and to infer the developmental history of specific subtypes of adult hypothalamic neurons. Finally, we demonstrate how the HyDD can be used to comprehensively analyze developmental mutants that generate complex phenotypes that would be difficult to characterize with traditional histology-based approaches, and in the process identify *Nkx2-1* as a negative regulator of prethalamic identity.

This study provides a reference atlas for future studies of hypothalamic development. It also identifies pathways by which gene regulatory networks that control cell identity can be targeted to analyze the functional role of individual hypothalamic neuronal subtypes.

## Results

### Comprehensive profiling of entire course of hypothalamus development

To profile changes in gene expression across the full course of mouse hypothalamic development, we processed 12 time points ranging from embryonic day (E)10 to postnatal day (P)45. For E10-E16, both prethalamus and hypothalamus were collected, whereas for E18-P45, only hypothalamus was profiled (Fig. 1a). In total, 129,151 cells were profiled (Fig. S1a,b). Using molecular markers of known hypothalamic regions and cell types^2^, we were able to annotate all major hypothalamic and adjacent brain regions, and major cell types at each individual age (Fig. S1c,d). Aggregation of the entire dataset with UMAP shows a clear developmental progression from neuroepithelial cells at E10, hypothalamic patterning between E11 and E13, and to the detection of major cell types in the mature hypothalamus (Fig. S2a). Roughly similar detection of expressed genes and total mRNAs were observed at each time point (Fig. S2b,c).

**Figure 1.**
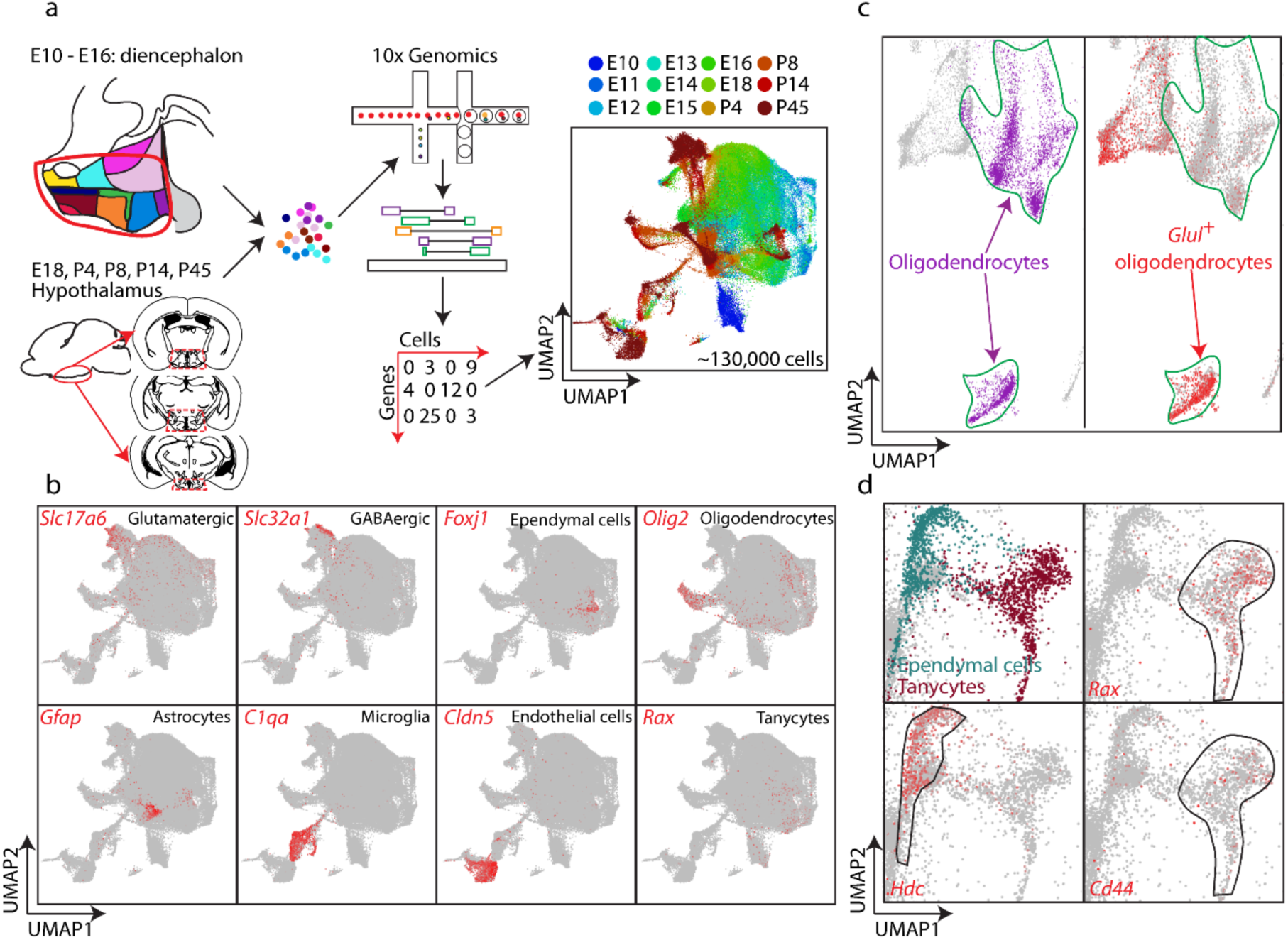
Overview of generation of the hypothalamus scRNA-Seq dataset. (a) Schematic diagram showing overall experimental strategy. 12 time points of developing diencephalon including the prethalamus and hypothalamus (between E10 and E16), and hypothalamus (between E18 and P45) were profiled using the 10x Genomics Chromium system. Distribution of individual ages (blue = younger time point, red = older time point) is shown in the UMAP plot. (b) UMAP plot showing distribution of major cell types (red) of the hypothalamus at the terminal neuronal branch. (c) UMAP plot (higher power view) showing two separate populations of oligodendrocytes, *Glul*-positive and *Glul*-negative oligodendrocytes in the hypothalamus. (d) UMAP plot (higher power view) showing diverging developmental trajectories giving rise to leading to ependymal cells (green, *Hdc*) and tanycytes (brown, *Rax, Cd44*).

Trajectories leading from neuronal progenitors to mature glutamatergic and GABAergic neurons, glia, ependymal cells and tanycytes were observed, as were separate clusters representing hematopoietic, microglial and endothelial cells (Fig. 1b). A separate cluster of oligodendrocytes was observed from P4 onwards, marked by a high and selective expression of *Glul* (Fig. 1c). Oligodendrocytes from posterior brain regions have been recently reported to selectively express high levels of *Glul*^*29*^, suggesting that these cells may migrate into the early postnatal hypothalamus from a more posterior location. Abundant expression of hypothalamic neuropeptides including *Pomc, Agrp, Ghrh, Sst, Gal, Hcrt*, and *Pmch* were observed in the neuronal cluster (Fig. S3).

Glial cells of the hypothalamus have been shown to play critical and tissue-specific roles in regulation of osmolarity^30^, circadian rhythm^31^, metabolism^32^ and neurogenesis^33^. To better understand the molecular mechanisms controlling the specification and differentiation of hypothalamic glia, each glial population was re-clustered and examined separately.

Cells that were identified as part of the oligodendrocyte maturation trajectory, and hence that share a similar molecular history, were re-clustered as previously described^13,34^, and genes that demarcate each stage of oligodendrocyte development were identified (Fig. S4a-c, Table S1). To identify genes selectively enriched in hypothalamic oligodendrocytes, mature oligodendrocytes were directly compared to scRNA-Seq datasets from mature cortical oligodendrocytes. While *Pcsk1n* and *Cbx3* are highly enriched in hypothalamic, relative to cortical, oligodendrocytes (Fig. S4d-h), these genes are enriched in all hypothalamic glial cells, and are not specific to oligodendrocytes.

In contrast, we identified many genes that were both astrocyte-enriched relative to other glial cell types, and selectively expressed in hypothalamic, relative to cortical, astrocytes (Fig. S5a-c). These include higher expression of *Agt*, and a lower level of *Mfge8* in hypothalamic astrocytes, as previously reported^34^, along with newly identified hypothalamic-enriched genes such as *Marcks* and *Marcks1* (Fig. S5a-c), which are important regulators of protein kinase C-dependent calmodulin signaling^35,36^.

Analysis of the developmental trajectory connecting gliogenic progenitor cells and hypothalamic astrocytes identified transitional states between these two populations. Immature hypothalamic astrocytes co-express the mature astrocyte marker *Agt*, and *Rgcc*, a cell-cycle regulator that regulates Notch signaling^37,38^ (Fig. S5d-f, Table S2). Loss of expression of genes specific to gliogenic progenitors was observed in hypothalamic astrocytes and other glial populations after the second postnatal week (Fig. S5g). Up-regulation of Notch signaling pathway components was also observed, as previously reported for human astrocyte development *in vitro*^*39*^(Fig. S5h).

Analysis of developmental trajectories for individual hypothalamic cell types identified the age at which these cell types began to diverge in gene expression, and identified both known and candidate regulators of cell fate. This is clearly seen when comparing the development of two ventricular glial-like cell populations -- ependymal cells and tanycytes. These two classes of ventricular cells begin to diverge at E13, with differential expression of *Foxj1* and *Rax*, markers of ependymal cells and tanycytes, first detected at this age (Fig. 1d, Fig. S6). Pseudotime analysis using BEAM analysis identifies additional transcription factors that are candidates for controlling tanycyte and ependymal cell specification and differentiation (Fig. S6a,b, Table S3). Tanycytes are themselves heterogenous, and can be subdivided into alpha and beta subtypes based on both spatial location and molecular markers^13,40^. To determine whether transcription factors enriched in differentiating tanycytes might also control tanycyte subtype specification, we analyzed previously published scRNA-Seq data obtained from mature tanycytes^14^ (Fig. S6c, d), and identified multiple tanycyte subtype-specific transcription factors that are expressed during early stages of tanycyte differentiation, and hence are strong candidates for controlling tanycyte subtype specification.

### Identification of genes selectively expressed in different regions of the developing hypothalamus and prethalamus scRNA-Seq

We next investigated whether we could use this dataset to faithfully distinguish hypothalamic domains that are spatially distinct in the embryo. To do this, we re-clustered data from E11, E12, and E13, which correspond to the peak period of hypothalamic neurogenesis (Fig. 2a)^41^. Using previously identified region-specific markers as a reference^2^, we observed a clear segregation of spatially-distinct neuronal precursors and progenitors (Fig. 2b, S7). We were able to readily distinguish hypothalamic and adjacent cell populations including the prethalamus, discrete clusters for telencephalic structures such as preoptic area and medial ganglionic eminence, thalamic eminence, rim domain, and main body of the sensory thalamus, as well as the zona limitans intrathalamica (ZLI) at all three developmental ages (Fig. 2b, S7-S8).

**Figure 2.**
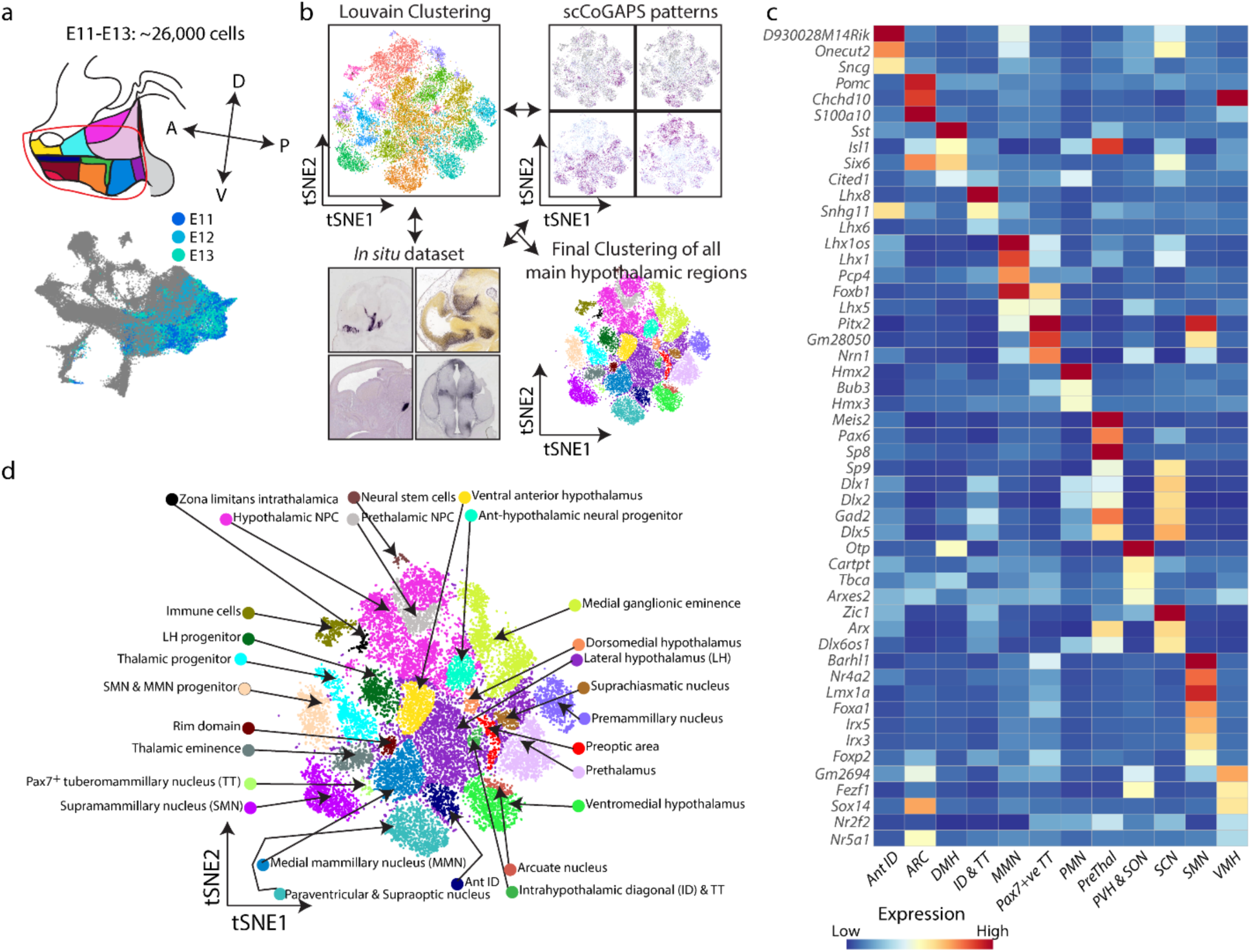
Specification of hypothalamic patterning during embryonic development. (a) Schematic diagram showing extraction of E11, E12 and E13 data to perform detailed analysis on hypothalamic patterning during development. (b) tSNE plot showing major subdivisions of the developing diencephalon (prethalamus and hypothalamus) and nearby regions to the developing diencephalon that are derived by cross-validation using the Louvain clustering algorithm, patterns from scCoGAPS and *in situ* analysis. (c) Heatmap showing a key subset of pattern-specific genes in major hypothalamic regions and prethalamus. (d) tSNE plot of E11-E13 developing diencephalon and adjacent regions.

Each of the previously reported major subdivisions of the developing hypothalamus^2^ were also identified, including postmitotic neuronal precursor cells of the paraventricular nucleus/supraoptic nucleus (PVN/SON), extrahypothalamic diagonal (ID) and tuberomammillary terminal (TT), ventromedial hypothalamus (VMH), arcuate nucleus (ARC), premammillary hypothalamus (PMN), mammillary nucleus (MMN), and supramammillary nucleus (SMN) (Fig. 2c,d, S7). In addition, several spatially distinct subtypes of mitotic hypothalamic progenitor cells were also observed, most notably cells that shared markers of both MMN and SMN (Fig. 2d, Table S4).

Multiple known and previously undescribed molecular markers, including many transcription factors, were identified for each of these regions (Fig. 2c, Table S4). While some of these markers are shared among multiple regions in the hypothalamus and other forebrain regions, others are highly specific and non-overlapping. We identified clear separation between mitotic neural progenitors and postmitotic neural precursors (Fig. 2d, S7).

This analysis was able to efficiently identify gene expression patterns that were restricted to specific spatial domains and subdomains of the developing hypothalamus and prethalamus, confirming and extending our previous findings^2^. However, this approach does not allow us to identify genes with more complex expression patterns, but which nevertheless may play important roles in regulating hypothalamic neurogenesis. To address this, we used scCoGAPS, a non-negative matrix factorization tool that allows unbiased identification of patterns of co-expressed genes^42^ (Fig. S8). Using this method, we identified patterns that not only matched key spatial subdivisions of the hypothalamus and prethalamus, but also patterns that labeled discrete subsets of hypothalamic progenitors (Table S5). These include hypothalamic neural precursor cells (NPC) that likely correspond to radial glia (*Fabp7, Slc1a3*), as well as neurogenic progenitors (*Pitx2, Nhlh1, Nhlh2*). Most strikingly, we observed multiple patterns that selectively label neurogenic progenitors that are located along the borders of the hypothalamus and prethalamus with both the telencephalon and ZLI. Genes that drive this pattern include *Neurog2, Lhx5*, and *Nhlh2* (Table S5). Although these expression patterns have been previously reported^2^, the fate of these border cells is unknown.

Due to the high complexity of the hypothalamic clusters observed in both two- and three-dimensional analysis, it is difficult to comprehensively visualize region-specific differences in gene expression. To improve visualization of these data, we generated a heatmap for major pattern marker genes that corresponds to the two-dimensional sagittal plane, capturing the main spatial subdivisions of the developing hypothalamus and adjacent brain regions (Fig. S9).

This analysis also identified clusters that correspond to three hypothalamic regions that had not been described in previous work^2^, including two populations of excitatory neurons. The first of these regions is found in the dorsomedial hypothalamus, and is marked by expression of *Sst, Cited1, Otp* and *Six6* (Fig. 2c). The second region is found in the TT/PMN region, and expresses *Pax7* (Table. S4). The third region is found in the lateral hypothalamus (LH), and consists of a diverse collection of subtypes of neuronal precursors. This LH cluster consists primarily of glutamatergic neurons, with a small subpopulation of GABAergic neurons (Fig. S10, Table S6). The glutamatergic population includes a discrete subcluster of *Lhx9*-positive neurons, which marks precursors of hypocretin neurons^2,43,44^. Cells within this LH cluster express multiple transcription factors that are also selectively expressed in other hypothalamic regions, including the VMH, PMN, MMN and ID.

Clustering of previously characterized spatial domains also identified discrete subclusters that express common sets of genes. This is clearly seen in the PVN/SON cluster (Fig. S11, Table S6). Selective expression of *Onecut2, Cartpt* and *Zic1* characterizes a ventrolateral domain that, based on its position, likely corresponds to the developing SON (Fig. S11). This same approach can be readily applied to other forebrain regions. We have previously identified molecular markers that both identify discrete spatial domains within the prethalamus, which gives rise to structures such as the thalamic reticular nucleus and ventral lateral geniculate nucleus^2,45,46^, and investigated whether these regions could be identified using scRNA-Seq data.

Sub-clustering of prethalamic cells allowed us to detect these and other spatial subdivisions within the prethalamus. We observed partially overlapping domains of expression of the transcription factors *Sp8* and *Sp9* (Fig. S12, Table S6), which play critical roles in the development of telencephalic interneurons^47^. *In situ* hybridization analysis revealed enriched expression of *Sp8* and *Lhx1* in anterior prethalamus and ID, while *Sp9* and *Prox1* were enriched in posterior prethalamus (Fig. S12). *Zic1* and *Ebf*1 also marked distinct, but partially overlapping spatial domains in prethalamus (Fig. S12).

Sub-clustering of the VMH allowed us to detect two distinct clusters, which corresponded to separate anterior and posterior domains of gene expression (Fig. S13, Table S6). A clear distinction between these anterior and posterior domains was detected until E16, both spatially and at the molecular level (Fig. S13). These two clusters had begun to spatially intermingle, yet the molecular distinction still remained, possibly reflecting local tangential cell migration within the VMH.

By combining our analysis of both the molecular markers of differentiation of major hypothalamic cell types and the selective markers of the different spatial domains of the developing hypothalamus and prethalamus, we have compiled a reference set of molecular markers that will be useful for further functional studies. We have designated this integrated and annotated scRNA-Seq dataset as a HyDD, or the *Hy*pothalamus *D*evelopmental *D*atabase.

### HyDD can rapidly annotate existing hypothalamus scRNA-Seq dataset and identifies developmental origins of VMH neurons

To demonstrate the broad usefulness of the HyDD, we first annotated a previously published scRNA-Seq dataset obtained through selective dissection of *Pomc-EGFP*-expressing cells from E15.5 hypothalamus using regional and cell type-specific markers from the HyDD^48^. In this study, while one cluster (cluster 0) was previously identified as the developing ARC, the remaining clusters were not annotated owing to the lack of well defined regional and cell type-specific markers to resolve spatial location of these clusters. Using markers obtained from the HyDD to train the dataset, we were able to annotate all but two clusters, representing cells from multiple hypothalamic regions, including VMH, PMH, anterior ID, DMH, SCN, and ARC. Some clusters were composed of cells from multiple hypothalamic regions, which may explain some of the previous difficulties in annotating these cells (Fig. 3a,b). Two unannotated clusters appear to reflect contamination from the habenula and pituitary that occurred during dissection (Fig. S14). A subset of the neurons in the ARC cluster share molecular markers of neural precursors in the PMN and DMH, implying that these cells may have migrated to the ARC from these regions (Fig. S14).

**Figure 3.**
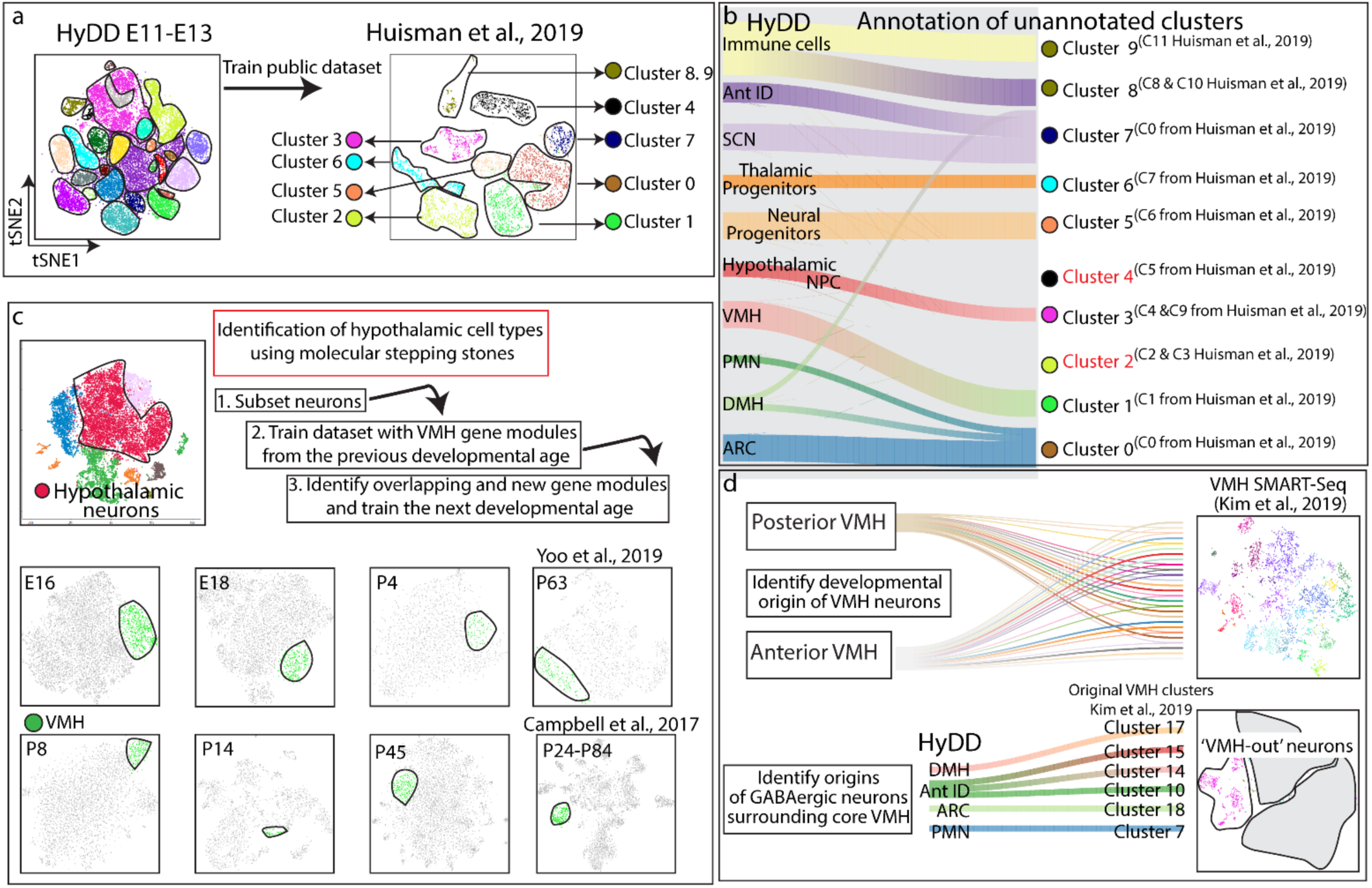
Utilizing HyDD to infer the identity and origin of individual cell types in the developing and adult hypothalamus. (a) The HyDD dataset was used to train a previously published scRNA-Seq on E15.5 hypothalamus obtained through selective dissection of *Pomc-EGFP*-expressing cells^48^. (b) Alluvial plot showing HyDD clusters (left) matched to clusters from Huisman et al., 2019^48^ (right). Note that 2 clusters (clusters 2 and 4) from Huisman et al., 2019^48^ do not match the HyDD dataset. (c) Using the molecular stepping stone approach to identify VMH neurons (green) across the entire developmental ages by identification of shared sets of gene modules that can demarcate the VMH across the entire hypothalamus scRNA-Seq dataset. (d) HyDD dataset is used to identify the developmental origins of previously annotated subtypes of glutamatergic neurons of the core VMH^51^ (top), and to identify the developmental origins of GABAergic neurons surrounding the core VMH (bottom).

We also identified a cluster that closely resembled hypothalamic NPC (Fig. S15), but which also co-expressed astrocyte-, ependymal and/or tanycyte-specific marker genes. Gene sets enriched in this cluster were then projected into the entire hypothalamus scRNA-Seq dataset (E10-P45), and glial populations including immature glial cells were enriched with these gene sets. This same gene expression pattern was found to be enriched in a subset of hypothalamic NPC that were detected from E11 onwards, and which may represent NPC that are competent to generate glia (Fig. S15). Many of these same genes are also expressed in the late-stage retinal progenitor cells, from the age at which they become competent to give rise to tanycyte-like Mũller glial cells^28^.

Since HyDD contains a nearly uninterrupted temporal profile of changes in gene expression during the process of cell specification and differentiation, HyDD can also be used to infer the developmental origins of fully mature hypothalamic neurons. However, identifying the precise spatial location of individual cell types from hypothalamus scRNA-Seq data based on specific molecular markers alone is bioinformatically challenging, due to the extreme tissue complexity. This is the case even when scRNA-Seq data has been generated with micro-dissected or flow-sorted cells from pre-defined hypothalamic regions. Most informative region-specific markers are strongly expressed early in development, but are either not expressed or show substantially different expression levels at later developmental ages^49^. Postmitotic hypothalamic neural precursors also undergo a considerable amount of tangential migration and dispersion, making it even harder to directly identify gene regulatory networks that control the specification of individual hypothalamic cell types^50^.

To identify the developmental origin of individual hypothalamic cell types, it is critical that overlapping sets of markers be identified that selectively label each stage of cell differentiation, in a manner analogous to molecular stepping stones, so that the developmental history of each cell type can be reconstructed. As a proof of principle for this approach, we identified gene sets that identify VMH cells at early stages of hypothalamic development (Fig. 3C), when region-specific molecular markers are robustly expressed. Gene sets specific to discrete spatial domains were then used to train the following developmental age to find VMH cells and new VMH-enriched genes were identified. This process was repeated for each successive developmental age. These VMH-enriched genes have varying levels of expression and specificity across the full course of the hypothalamus development (Fig. S16).

We next used the HyDD to identify the developmental origin of major VMH neuronal subtypes. Recent scRNA-Seq of the adult VMH identified multiple clusters of both core glutamatergic VMH neurons and of GABAergic neurons surrounding the core VMH (VMH-out)^51^. We sought to identify the developmental origins of both classes of VMH neurons. We first found that GABAergic neurons of VMH-out originated from four distinct regions of the developing hypothalamus - ARC, DMH, Ant ID and PMN (Fig. 3d) -- with each VMH-out GABAergic cluster having a distinct developmental origin based on specific expression of regional markers. We likewise observed that different subsets of core glutamatergic VMH neurons arise from distinct anterior or posterior domains of the embryonic VMH (Fig. S13). Some of these clusters remain restricted to anterior or posterior regions of the adult VMH, as noted in the original study^51^(Fig. 3d, Fig. S17a,b). However, the majority of VMH neuronal subtypes originate from both anterior and posterior domains of the developing VMH (Fig. S17), and are distributed widely along the anterior-posterior axis of the adult VMH^51^. VMH neuronal subtypes may thus be two distinct developmental steps: an initial stage in which anterior and posterior identity is specified between E11 and E13, and a later stage that coincides with the initiation of local tangential migration that occurs from E16 onwards.

### HyDD allow rapid and comprehensive analysis of complex mutant phenotypes that alter hypothalamic and prethalamic patterning

HyDD provides both a high-resolution molecular atlas of the developing hypothalamus and prethalamus, and a useful resource to understand the developmental origin of adult hypothalamic neurons. We next sought to determine if HyDD could also be used to rapidly and comprehensively characterize mutants that regulate early stages of hypothalamic development and organization. As proof of concept, we performed scRNA-Seq analysis on E12.5 *Foxd1*^*CreGFP/+*^;*Ctnnb1*^*ex3/+*^ mice, in which a constitutively active form of beta-catenin is overexpressed in *Foxd1*-positive hypothalamic and prethalamic progenitors, leading to activation of canonical Wnt signaling in these cells and their descendants^20^. The same analysis was also with *Foxd1*^*CreGFP/+*^ littermate controls. These mice show broad activation of the canonical Wnt pathway effector *Lef1*, a hyperplastic ventricular zone, and with the exception of a handful of posterior hypothalamic markers, show the loss of most regional markers in the hypothalamus and prethalamus^20^.

ScRNA-Seq analysis of control and mutant animals at E12.5 reveals several mutant-specific cell clusters (Fig. S18a). Using the HyDD to annotate both control and mutant data, we identified changes in gene expression and cell composition that match previously reported findings (Fig. 4a), where we observed a substantial increase in undifferentiated NPC, along with a corresponding reduction in the number of cells expressing markers of hypothalamic and prethalamic neuronal precursors (Fig. 4b, S18). In particular, strong loss of markers shared by both hypothalamus and prethalamus, such as *Arx, Isl1* and *Gad1/2* (Fig. S19, Table S7) was observed. We also identified two cell clusters that are found exclusively in mutant mice, both of which express NPC markers, and also highly express both *Lef1* and negative regulators of canonical Wnt signaling such as *Dkk1, Wif1* and *Axin2* (Fig. S18e).One of these clusters is strongly enriched for G2/M phase markers such *Ube2c, Rrm2*, and *Ccnb1* (Fig. 4b). Flow cytometry data also demonstrated a substantially higher fraction of NPCs in G2/M phase in mutant mice (Fig. S18f), as has been previously reported in non-neuronal cells that show high levels of canonical Wnt signaling^52^. This finding explains the previous observation that, although a massive increase in the number of NPC cells is seen in these mutants, only a modest increase is observed in EdU labeling, which labels S-phase NPC^20^. This demonstrates the power of using scRNA-Seq in conjunction with the HyDD to analyze developmental phenotypes, in a manner that is far more rapid and comprehensive than conventional histological techniques.

**Figure 4.**
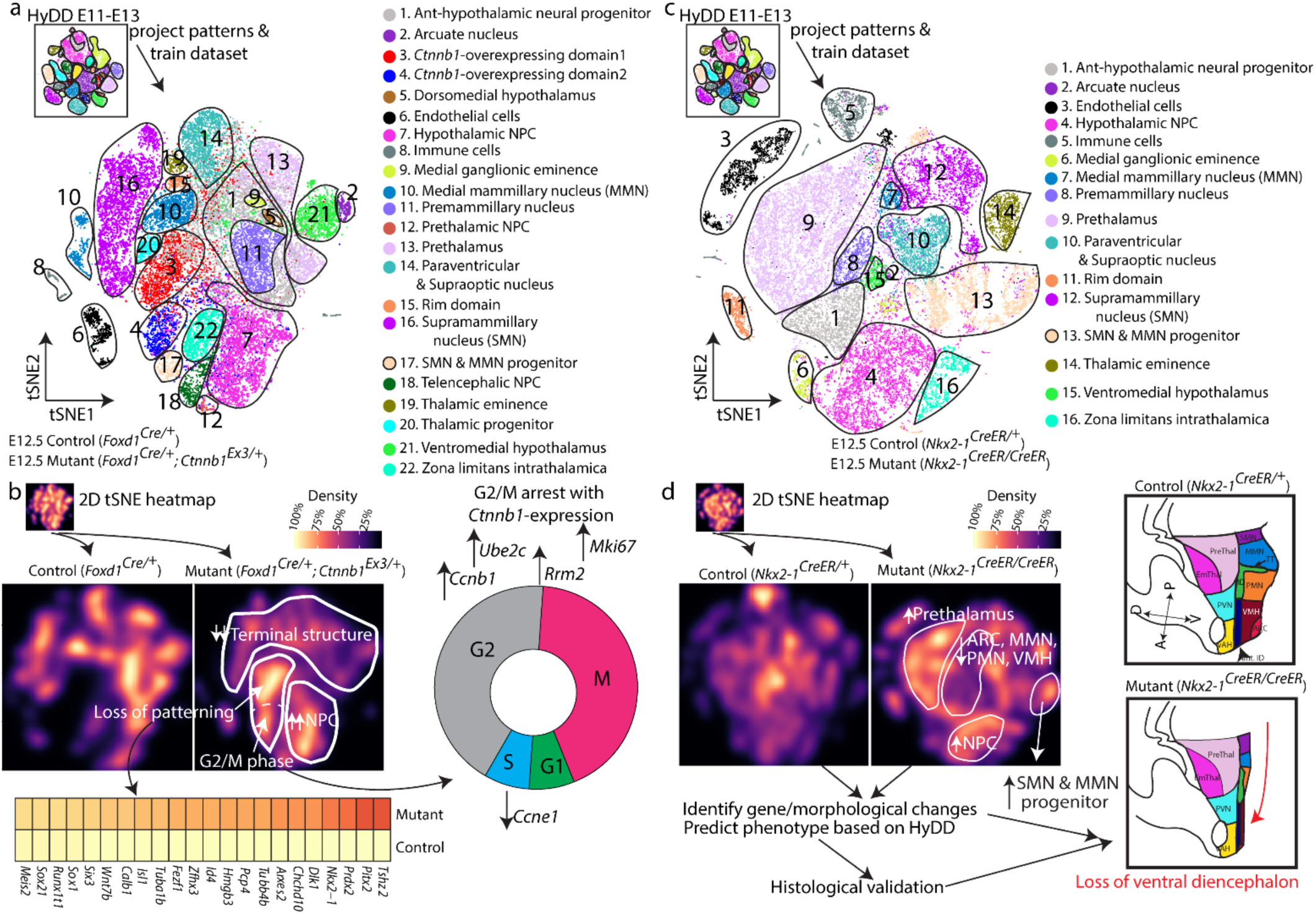
scRNA-Seq can be used to screen mutant phenotype in the hypothalamus. (a) tSNE showing distribution of diencephalic and nearby brain structures in control (*Foxd1*^*Cre/+*^) and mice selectively overexpressing a constitutively active mutant form of *Ctnnb1* in neuroepithelial cells of the hypothalamus and prethalamus (*Foxd1*^*Cre/+*^;*Ctnnb1*^*Ex3/+*^) with clusters obtained by training the dataset with HyDD markers. Note distinct clusters (clusters 3 and 4) that are only observed in mutant samples)^20^. (b) tSNE heatmap showing distribution of individual cluster between control (*Foxd1*^*Cre/+*^) and constitutively active *Ctnnb1* mutants (*Foxd1*^*Cre/+*^;*Ctnnb1*^*Ex3/+*^)(top left) and heatmap showing that mutant-specific clusters display all pattern-specific markers that are expressed across the developing diencephalon (bottom left), with a high level of G2/M phase markers in the mutant-specific cluster (right). (c) tSNE showing the distribution of hypothalamic, prethalamus and adjacent brain structures of control (*Nkx2-1*^*CreER/+*^) and *Nkx2-1* mutant line (*Nkx2-1*^*CreER/ CreER*^) with clusters obtained by training the dataset with HyDD markers. (d) tSNE heatmap showing distribution of individual clusters between control (*Nkx2-1*^*CreER/+*^) and *Nkx2-1* mutant line (*Nkx2-1*^*CreER/ CreER*^)(left). Note the absence of ventral diencephalic structures (except the supramammillary nucleus), and the relative expansion of the prethalamus in *Nkx2-1* mutants (right).

We next used this same approach to characterize E12.5 *Nkx2-1*^*CreER/CreER*^ knock-in mice, which are homozygous for a null allele in the homeodomain transcription factor *Nkx2-1*^*53*^. *Nkx2-1* is broadly and selectively expressed in ventral hypothalamic progenitors, as well as in progenitors that give rise to telencephalic interneurons^54,55^. Loss of function of *Nkx2-1* leads to a substantial reduction in ventral hypothalamic structures by E18^56^, but a detailed molecular characterization of these mutants has not been conducted.

Analysis of *Nkx2-1*^*CreER/CreER*^ mutants and heterozygous littermate controls revealed changes in cluster densities in the mutant (Fig. S20). We observed a broad loss of markers specific to *Nkx2-1* positive ventral hypothalamic structures such as ARC, VMH, PMN, and MMN, but not the SMN (Fig. 4c,d, Fig. S21, Table S8, S9), with both the relative expression levels and the number of cells expressing these markers reduced. Both the width of the hypothalamic ventricular zone and the levels of EdU incorporation were reduced in *Nkx2-1*^*CreER/CreER*^ mice (Fig. S22). An increase in the fraction of cells expressing prethalamic markers was detected (Fig. 4d, Fig. S23), and increased Cre expression was also observed in these mice.

In contrast to controls, prethalamic cells in mutant mice expressed *Cre*, implying that ventral hypothalamic cells that normally express *Nkx2.1* may have acquired prethalamic identity (Fig. S22). To investigate this further, RNAscope probes against *Sp9, Meis2* and *Cre* were used to visualize the location of these *Cre*-positive prethalamic cells, and substantial co-localization of prethalamic markers and *Cre* expression was observed in the region normally occupied the by the ventral hypothalamus in controls (Fig. S22). This implies that *Nkx2-1* not only maintains the identity of ventral hypothalamic progenitors but also actively represses expression of molecular markers of prethalamic identity. *In situ* hybridization confirmed that there was an increase in the absolute size of the prethalamus and its proportion in the diencephalon (Fig. S23). An increase in the number of cells expressing markers of NPC in the SMN and MMN was also seen, while *Nkx2-1* negative hypothalamic regions such as the PVN/SON are unaffected (Fig. 4d, S24).

## Discussion

In this study, we use scRNA-Seq to develop a molecular atlas of the developing mouse hypothalamus, with a particular focus on stages when hypothalamic patterning and neurogenesis are regulated. This dataset identifies genes that are selectively expressed during the differentiation of major neuronal and non-neuronal hypothalamic cell types, and accurately delineates spatial subdivisions present in the early stages of development of both the hypothalamus and the adjacent prethalamus. It also identifies many previously uncharacterized transcription factors and other genes that are excellent candidates for controlling regional patterning and specification of individual hypothalamic cell types. Combining functional analysis of these genes with the new selective markers of hypothalamic regions and immature hypothalamic cell types identified in this study has the promise to greatly expand our knowledge of hypothalamic development and organization.

The integrated dataset presented here provides three specific features that are critical for studying the formation and function of the hypothalamus. First, it makes it straightforward to unambiguously annotate major cell types at all stages of hypothalamic development. Second, it makes it possible in many cases to infer the developmental histories of hypothalamic cells in both the developing and mature hypothalamus. Third, it allows rapid and accurate phenotyping of mutants that show broad effects on hypothalamic patterning, neurogenesis and differentiation, with which we were able to validate our findings using traditional histological analysis. Despite the availability of highly specific molecular markers for the major spatial subdivisions of the hypothalamus, the highly complex and temporally dynamic anatomy of this brain region makes analysis of mutant phenotypes slow and complex. Previously, it has taken up to several years of full-time labor to obtain detailed characterization of individual mutant lines. The HyDD dataset allows these analyses to be conducted far more rapidly, efficiently, and comprehensively.

Our scRNA-Seq characterization of *Nkx2-1*-deficient mice identifies an unexpected developmental connection between the hypothalamus and prethalamus, where *Nkx2-1* can potentially act as both a positive regulator of ventral hypothalamic identity while simultaneously repressing prethalamic identity. This result is not predicted by the current prosomeric model for forebrain organization^57,58^, and raises questions about the early development and patterning of these structures. Previous models of hypothalamic development and organization were constructed using very sparse datasets - typically single color in situ hybridization of a limited number of genes at a small number of time points. The much richer datasets provided by scRNA-Seq, and interpreted using the HyDD data, offer a far more powerful resource for constructing these models..

## Contribution

DWK, SB designed experiments. DWK, PWW, ZQW, BTI, SL, LJ, HW performed experiments. DWK, PWW, ZQW, SL, CS analyzed data. All authors contributed to writing the paper

## Acknowledgements

This work was supported by a grant from the NIH (DK108230) to S.B. We thank Transcriptomics and Deep Sequencing Core at Johns Hopkins for sequencing all scRNA-Seq libraries, Ross Flow Cytometry Core (Johns Hopkins) for FACS analysis, and Microscope facility (Johns Hopkins MICFAC, supported by the award number S10OD018118). We thank M. Placzek, E. Newman, J. Nathans, A. Kolodkin, W. Yap, and members of the Blackshaw lab for comments on the manuscript.

## Data availability

All scRNA-Seq data are available on GEO, GSE132355. Data can be viewed at https://proteinpaint.stjude.org/F/mm10/example.scrna.html.

