## Supplemental text and Figures S1-S24 for "The cellular and molecular landscape of hypothalamic patterning and differentiation"

**List of Supplementary Materials**

### Materials and Methods:

#### Mice

All experimental animal procedures were approved by the Johns Hopkins University Institutional Animal Care and Use Committee. All mice were housed in a climate-controlled facility (14-hour dark and 10-hour light cycle) with *ad libitum* access to food and water. Time-mated CD1 or C57BL/6J mice were ordered from Charles River Laboratories to collect mice embryos at E10 (CD1), E11 (CD1), E12 (CD1), E13 (CD1), E14 (CD1), E15 (CD1), E16 (1 CD1 and 1 C56BL/6J), and E18 (CD1). Time-mated CD1 mice were ordered to collect pups at P4, P8 and P14. C57BL/6J mice were used for P45 samples.

*Nkx2-1*$^{CreERT2}$ knock-in[1](JAX #014552), *Foxd1*$^{CreGFP}$ knock-in[2](JAX #012463), *Ctnnb1*$^{ex3/ex3}$ [3] were used for single-cell phenotyping studies. Ctnnb1$^{ex3/ex3}$ mice were crossed with *Foxd1*$^{CreGFP/+}$ to generate *Foxd1*$^{CreGFP/+}$ or *Foxd1*$^{CreGFP/+}$;*Ctnnb1*$^{ex3/+}$. *Nkx2-1*$^{CreERT2}$ was C56BL/6J background, and *Foxd1*$^{CreGFP/+}$ knock-in, *Ctnnb1*$^{ex3/ex3}$ were C57BL/6 and CD1 mixed background. Mice were time-mated during their estrous cycle and vagial plugs were observed to detect successful mating. E12.5 embryos were collected for generating scRNA-Seq dataset.

#### Dissection and cell dissociation

Embryos or postnatal mice were collected and dissociated using a previously published protocol[4]. Embryos were collected using Hibernate-E media (Thermo Fisher Scientific) with 2% B-27 supplement (Thermo Fisher Scientific) and GlutaMAX supplement (0.5 mM final, Thermo Fisher Scientific). A small incision was made dorsal to the lower jaw to expose the ventral portion of the brain. For samples collected between E10 and E16, tissue residing posterior to the medial ganglionic eminence and anterior to the midbrain and sensory thalamus was dissected to collect both developing prethalamus and hypothalamus. Prethalamus was excluded from samples aged E18 and older, with only hypothalamus collected, as previously described[5]. Exclusion of medial ganglionic eminence (anterior to the hypothalamus) and structures posterior to the supramammillary nucleus ensured that the equivalent diencephalon area (hypothalamus and prethalamus) was always included. Between 8 and 12 embryos of either sex were collected for each embryonic time point and pooled for scRNA-Seq dataset.

Postnatal mice were collected using Hibernate-A media with 2% B-27 and GlutaMAX (0.5 mM final), and the region that is posterior to the optic chiasm (Bregma -0.58 mm) and anterior to the hypothalamus-midbrain border (Bregma 2.54 mm) were collected. Eight pups (4 male and 4 female) were collected for P4, P8, and P14 dataset, and 3 male mice were pooled for P45 dataset. E10, E12, and E15; E11 and E13; E14, E16, and P45; E18, P4, and P14 were each generated on the same day. Consecutive developmental ages were not collected and processed on the same day and this ensured that trajectories identified using tSNE and UMAP are not determined by batch-effect but by developmental stages.

For single-cell phenotyping studies, E12.5 time-mated embryos were collected and placed in a buffer mentioned above on ice. Tail-tips were collected and rapidly genotyped using GeneAmp Fast PCR master mix (Thermo Fisher Scientific). Both control and mutant groups were collected on the same day, and E12.5 embryos were pooled from 3 different dams. Between 6 and 8 embryos were collected to generate individual scRNA-Seq libraries.

Following dissection, tissues were dissociated in papain (Worthington Biochemical) as previously described in calcium-free Hibernate media [4]. Tissue debris were removed using OptiPrep density gradient media (Sigma-Aldrich) in postnatal mice following cell dissociation. Numbers of viable cells were counted manually via haemocytometer with Trypan Blue staining and cross-checked with an automated cell counter, and cell concentration was adjusted following the manufacturer's protocol of 10x Genomics.

**ScRNA-Seq library generation and data processing**

Suspended cells were loaded into 10x Genomics Chromium Single Cell System (10x Genomics), and libraries were generated using v1 (1 library) and V2 chemistry with manufacturer's instructions. Libraries were sequenced on Illumina MiSeq (1 library) and NextSeq500 high-output (400 million reads). Sequencing data were first pre-processed through the Cell Ranger pipeline (10x Genomics, Cellranger count v2.0.2) with default parameters (expect-cells set to number of cells added to 10x system), aligned to mm10 genome (refdata-cellranger-mm10-1.2.0), and matrix files were used for subsequent bioinformatic analysis.

**Data analysis**

For analysis of the entire hypothalamic dataset, Seurat V2[4,6] and Scanpy[7] were used to process matrix files to keep all cells with at least 200 detected genes. Data-set were normalised to transcript copies per 10,000, and log-normalised to reduce inter-batch effect and adjust for individual variation. For the entire hypothalamic dataset, the top 4000 highly variable genes were used for Principal Component Analysis (PCA), and 100 PCA variables were used for either tSNE, to preserve local structures and to visualize complex clusters in two-dimensional space without minimum influences from global structures (developmental stages), or UMAP, to preserve global distances to better visualise changes across developmental stages[8]. Log-normalised values with xyz coordinates from UMAP or xy coordinates from tSNE were extracted, and plots were drawn using a custom-written script.

Individual ages were initially clustered using the Louvain clustering algorithm provided in Scanpy with default parameters. Individual developmental ages were highlighted in the UMAP dataset, and individual cell types (or hypothalamic regions) were projected intoUMAP to determine segregation of cell types (or regional markers) in the dataset regardless of developmental ages (https://proteinpaint.stjude.org/F/mm10/example.scrna.html).

To identify region-specific differences between mature oligodendrocytes or astrocytes of the hypothalamus and other brain regions, previously published cortical scRNA-Seq[9,10] were used to identify differential gene expression, using age as the variance with default parameters (first-pass gene lists)[11]. Published hypothalamic datasets[10,12–14] were then used to compare to these cortical scRNA-Seq datasets using the identified genes (second-pass gene lists). Identified differential genes were then validated by matching the Allen Brain *in situ* atlas data using co-coframer[15] to validate spatial expression of these differential genes, as some observed differences could reflect batch effects resulting from scRNA-Seq library preparations from different laboratories (third-pass gene lists).

To identify the developmental origins of ependymal cells and subtypes of tanycytes, transcription factors that are highly expressed near the midpoint of the pseudotime branch -- where cells are not full mature tanycytes but no longer gliogenic progenitors -- were extracted and clustered to scRNA-Seq data obtained from adult tanycytes[13,16] using Garnett[17].

For analysis of hypothalamic patterning and generation of the developmental database HyDD (*<u>Hy</u>*pothalamus *<u>D</u>*evelopmental *<u>D</u>*atabase), the E11, E12, and E13 datasets were used to perform a detailed analysis of hypothalamic patterning[5], and processed as described above using the top 2000 highly variable genes with 30 PCA variables. Initial clustering was conducted using the Louvain clustering algorithm in Scanpy with 0.6 resolution[18], and individual clusters (initially divided based on rough anatomical locations and by cell cycle status) were further subdivided to capture all the main subdivisions of the developing hypothalamus, prethalamus, and adjacent structures. The initial clustering results were then cross-referenced to patterns from scCoGAPS, which were superior in capturing changes in gene expression over time, and selective markers of small sub-regions of the developing hypothalamus and prethalamus.

Cross-referencing between these two pipelines allowed us to identify all the sub-regions of the developing hypothalamus, prethalamus, and adjacent structures. Following sub-divisions, enriched genes (top 50 highly-enriched genes in an individual cluster) in the individual cluster were extracted and cross-referenced back to our previous work[5], *in situ* hybridization (ISH) validation, Allen Brain *in situ* atlas[15,19] and GenePaint to further validate our cluster assignments[15,19], as well as identifying pattern-specific markers. The final main clusters were subsetted and reclustered to identify additional subdivisions within the major hypothalamic and prethalamic regions. This was repeated until the child clusters could not generate any further clusters that had differential expression of transcription factors or no visible spatial distinction between markers under the Allen Brain *in situ* atlas.

For clustering previously published datasets[20], data was processed as previously described in the original paper. The HyDD dataset was used as the reference point to train the dataset using Garnett, using the top 10 most selective markers identified from individual HyDD clusters, and identified clusters that were aligned to our dataset. For unannotated or poorly annotated clusters (which comprised less than 5% of all individual cells trained by our dataset), markers were identified and checked using the Allen Brain *in situ* atlas. These clusters were located outside of developing hypothalamus, in the habenula and pituitary, further validating the accuracy of the molecular markers identified in this study. An alluvial plot was then generated based on the percentage of clusters that were trained using our HyDD dataset. The opposite approach was taken to cross-validate HyDD annotation and to identify NPC that express high levels of gliogenic gene sets.

To identify VMH cells across the entire course of hypothalamic development using the molecular stepping stone approach, gene sets identified as labeling the VMH from the E11-E13 datasets used to generate HyDD (markers that are significantly expressed in the VMH but not in ARC, Table S4), were used to train the next developmental age (E14) and identify VMH from E14 scRNA-Seq dataset using Garnett[17]. Following the identification of the VMH, scCoGAPS was used to identify a VMH-specific gene expression pattern (i.e. the set of expressed genes that can selectively demarcate the VMH), and identified new and overlapping gene sets of VMH cells. These gene sets were then used to train the next developmental age (E15), and this process was repeated to the oldest age (P45) in our scRNA-Seq dataset. Genes that could identify the VMH across at least 3 developmental ages, or at least 2 developmental ages if the gene in question could also selectively specific cell clusters in the adult VMH, were selected as the final VMH gene sets (Fig. S16). This process was necessary to identify cells comprising specific hypothalamic nuclei, since at older ages, genes that are highly informative at providing information about spatial localization within the hypothalamus are often no longer highly expressed. This molecular stepping stone approach was validated by identification of glutamatergic VMH neurons using the public droplet-based scRNA-Seq dataset[13,16]. Spatial information of the identified clusters were validated by comparison to the Allen Brain *in situ* atlas data using co-coframer[15], as well as by matching to the published SMART-Seq data obtained from the adult VMH[21].

To identify the developmental origins of GABAergic neurons surrounding the core VMH (VMH-out), clusters obtained from public SMART-Seq data capturing only VMH[21], were generated to match the previously published clusters. VMH-out GABAergic cells were then trained using HyDD region-specific markers, as expression of many region-specific markers persisted into the adult stage (although usually at low levels),  and could readily be detected by SMART-Seq.

To identify the developmental origin of major cell clusters in the VMH, key gene sets from the anterior and posterior VMH were extracted from HyDD (Fig. S13, Table S6). These gene sets of the anterior and posterior VMH were validated in both our dataset at later developmental ages and in a published dataset using the molecular stepping stone approach[22]. Additional histological validation was conducted  using RNAscope (described below) and using the Allen Bran *in situ* atlas, anterior and posterior VMH gene sets were used to train VMH clusters. Some clusters clearly labeled either anterior or posterior domains of the embryonic VMH, and were also restricted to the corresponding region of the adult VMH[21], validating this approach.

For mutant phenotyping, both control and mutants of individual lines were first merged together using the above method. Given the complex phenotypes of these mutants, we used sets of region-specific markers obtained from HyDD, we trained each individual dataset using Garnett[17], which allowed us to faithfully cluster the mutant dataset, since the majority of pattern-specific markers were expressed in both genotypes, although often at different cellular expression levels. This approach allowed us to identify clusters that did not match annotated regions in HyDD, indicating the presence of mutant-specific clusters. Gene expression differences between control and mutants were compared between each identified region of the developing hypothalamus and prethalamus. For unidentified regions in mutant samples, gene expression was compared to all control regions. The percentage of regions occupied by cells of either genotype was compared as well. Both changes in pattern-specific markers and percentage of clusters occupied by each genotype reflect biological differences, but not changes in gene expression that resulted from variation in conditions occurred during dissection and library preparation. Top differential pattern-specific markers (mostly transcription factors) in regions that were differentially occupied between control and mutants were then selected for histological validation. xy coordinates from tSNE were used to estimate density (contour heatmap) of individual clusters between control and mutant groups.

Differential gene tests on the scRNA-Seq datasets were initially performed using Seurat V2 *FindAllMarkers* using MAST with default parameters on all expressed genes by using individual dataset (age or genotype) as a variance, and cross-referenced to Monocle2 VGAM likelihood ratio tests using age or genotype as the full model were then used in differential expression tests with default parameters.

**scCoGAPS analysis**

scCoGAPS, a Bayesian non-negative matrix factorization algorithm, was used to identify “patterns”, or  sets of co-expressed genes and their cellular expression levels, their weights in either the E11-E13 dataset or the individual/combined E10-P45 dataset, in order to identify cells that represent gliogenic progenitors or immature glia, or to identify VMH at individual developmental ages, using previously described parameters[23].

All genes were used to identify patterns for E11-E13 dataset, scCoGAPS patterns were projected into our tSNE plot and interpreted by using our prior knowledge of gene expression patterns in the developing hypothalamus. For the molecular stepping stone approach used to trace the developmental history of VMH neurons, or for identifying gliogenic progenitor populations. 200 HyDD cell or region-specific markers were used to annotate patterns, with the ‘patternMarkers’ function in order to identify robust pattern markers across developmental stages.

**E12 spatial mapping**

xy coordinates of E12 were drawn based on our previous work to capture all post-mitotic regions of the developing hypothalamus and prethalamus, average values of individual clusters were assigned to each XY coordinate, and 2D spatial representation was produced using the Python-based SpatialDE package [24,25].

**Cell cycle analysis**

Cell cycle data for NPCs from *Foxd1*$^{Cre/+}$;*Ctnnb1*$^{Ex3/+}$ mice overexpressing constitutively active *Ctnnb1* in hypothalamus and prethalamus, as well as age-matched controls, were analyzed using scran [20]. Additional cell cycle analysis was conducted by staining dissociated E12.5 control and mutant developing diencephalon with propidium iodide and analysed DNA content using LSR II (BD) and FlowJo (v10.6.1).

**Pseudotime analysis**

Monocle2[11] was used to perform pseudotime analysis to identify differences in gene expression in differentiating tanycytes and ependymal cells.

***In situ* hybridization (ISH)**

Chromogenic *in situ* hybridization was performed as previously described[5], except E12.5 or E13.5 embryos were fixed with 4% paraformaldehyde, sectioned at 25 um with either coronal or sagittal plane and treated with proteinase K for 5 min at room temperature.

Single-molecule fluorescence *in situ* hybridization was performed using RNAScope with probes targeting *Meis2*, *Sp8*, *Sp9*, *Pitx2*, *Nhlh2*, *Tcf7l2*, *Foxp2*, *Snca*, and *Cre* on E12.5 or E13.5 wild-type or *Nkx2-1*$^{CreERT2/CreERT2}$ embryos; and *Nhlh2*, *Foxp2*, *Snca* on E13.5, E16.5, E18.5, and P4 wild-type mice following the manufacturer’s protocol.

**EdU**

EdU (20 mg/ml) was injected 3 hours prior to collection into time-mated dams, and embryos were collected and processed, as previously described[5,26].

#### Immunostaining

Fixed embryos were processed for immunostaining with Pax6-antibody (1:200, AB2237, EMN Millipore). Sections were mounted with Vectamount (Vectorlabs) and imaged under Keyence BZ-X800 fluorescence microscope and Zeiss LSM 700 microscope.

a

| Age | Cells | 10x |
|---|---|---|
| E10 | 3927 | 10x V2 |
| E11 | 7171 | 10x V2 |
| E12 | 9310 | 10x V2 |
| E13 | 9677 | 10x V2 |
| E14 | 16543 | 10x V2 |
| E15-1 | 10867 | 10x V2 |
| E15-2 | 2698 | 10x V2 |
| E16-1 | 16792 | 10x V2 |
| E16-2 | 1134 | 10x V1 |
| E18 | 8865 | 10x V2 |
| P4 | 4521 | 10x V2 |
| P8 | 13677 | 10x V2 |
| P14 | 6944 | 10x V2 |
| P45-1 | 8084 | 10x V2 |
| P45-2 | 8941 | 10x V2 |
| **Sum** | **129151** | |

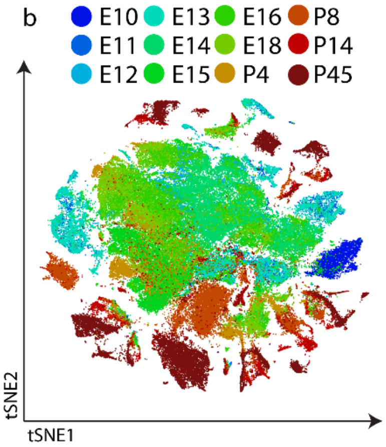


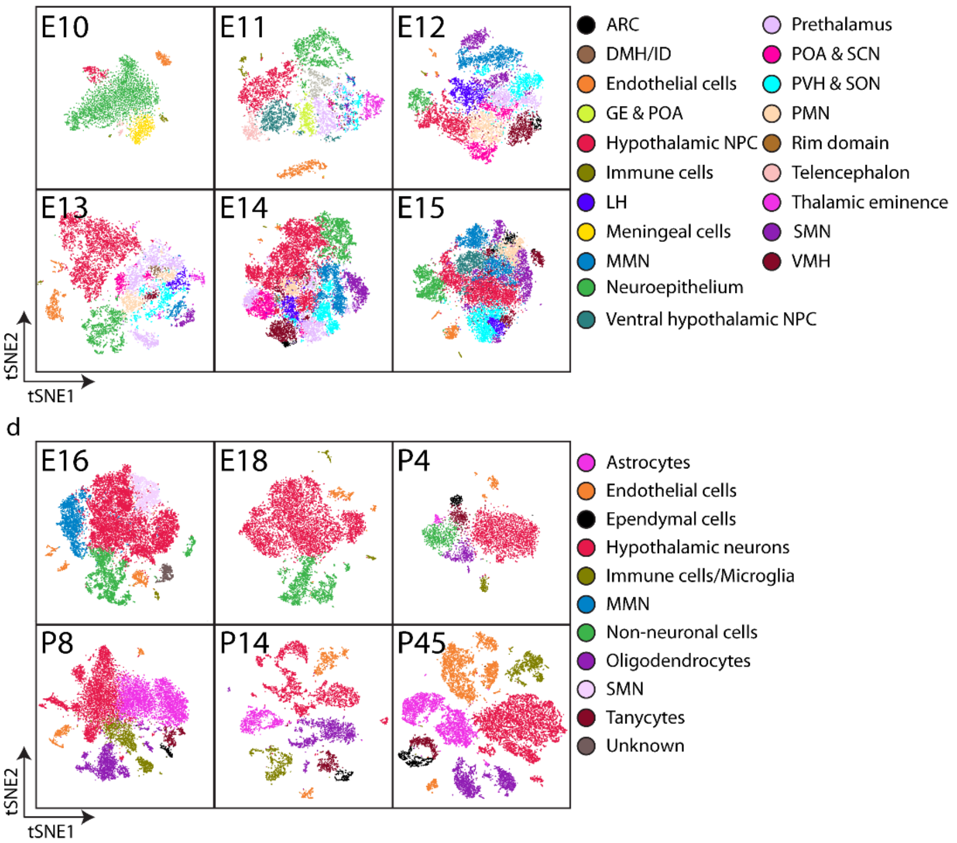


ExtFig1. Kim, et al.

**Extended Figure 1. Overview of individual time points collected for scRNA-Seq analysis.** (a) Table showing number of cells collected for scRNA-Seq dataset. (b) tSNE showing distribution of individual ages. (c) tSNE showing distribution of individual prethalamic and hypothalamic regions between E10 and E15. (d) tSNE showing distribution of individual major cell types of the hypothalamus between E16 and P45.

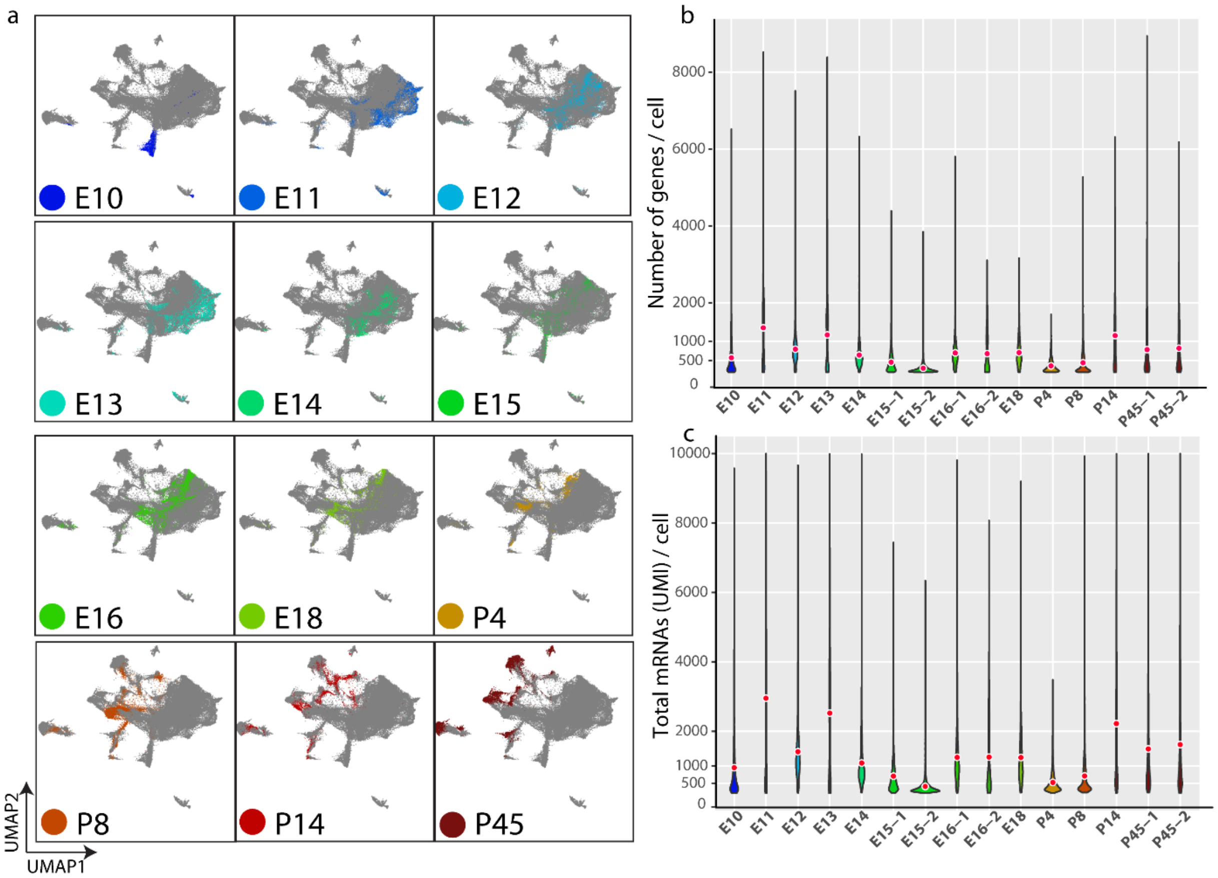


ExtFig2. Kim, et al.

**Extended Figure 2. Overview of individual ages using UMAP plotting.** (a) 3D UMAP plot (which is shown at a different angle than in Fig. 1 to better show the distribution of individual ages) showing distribution of individual ages collected for scRNA-Seq. Note that the UMAP plot shows developmental trajectories. (b) Violin plot showing distribution of mean (black dot) and of number of genes in individual scRNA-Seq libraries. (c) Violin plot showing distribution of mean (black dot) and number of total mRNAs (UMI) in individual scRNA-Seq libraries.

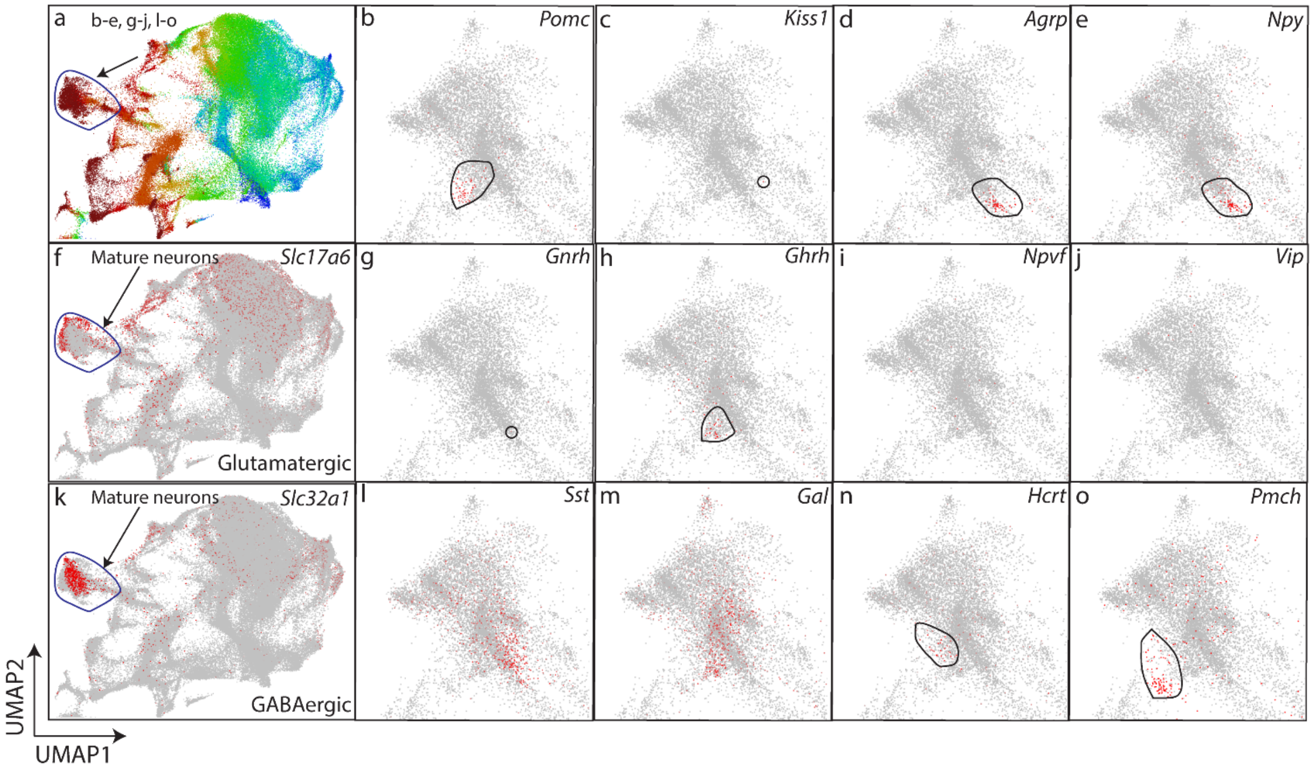


ExtFig3. Kim, et al.

**Extended Figure 3. Distribution of hypothalamic neuronal cell types.** (a) 3D UMAP plot (which is shown at a different angle and magnification than in Fig. 1 to highlight hypothalamic neurons) highlighting the terminal trajectory of hypothalamic neurons (black line). (b-e) UMAP plot focusing on hypothalamic neurons (magnified to focus on hypothalamic neurons) highlighting expression of *Pomc* (b), *Kiss1* (c), *Agrp* (d), *Npy* (e). (f) UMAP plot highlighting expression of *Slc17a6*. (g-j) UMAP plot highlighting expression of *Gnrh* (g), *Ghrh* (h), *Npvf* (i), *Vip* (j). (k) UMAP plot highlighting expression of *Slc32a1*. (l-o) UMAP plot highlighting expression of *Sst* (l), *Gal* (m), *Hcrt* (n), *Pmch* (o).

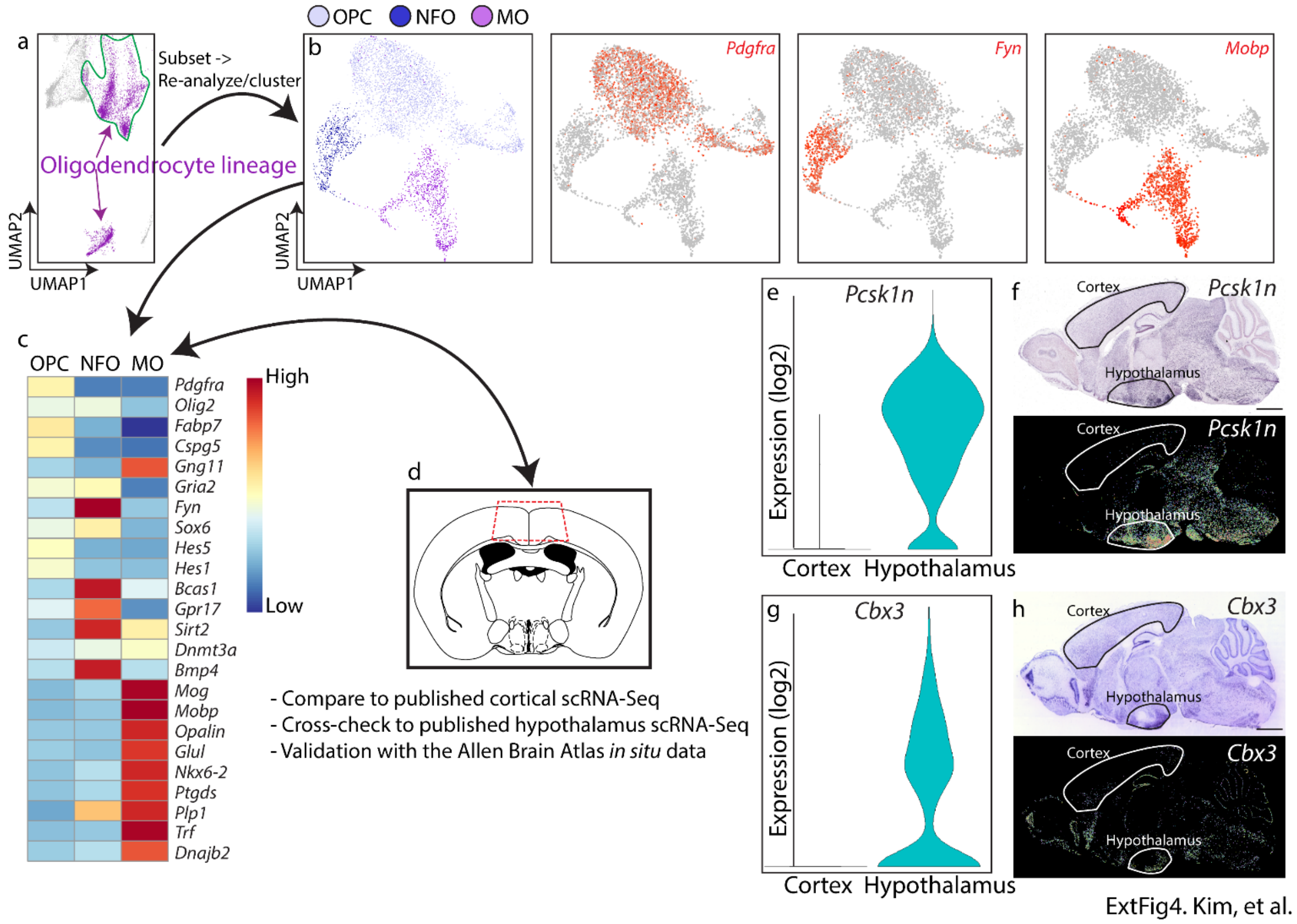


**Extended Figure 4. Dynamics of gene expression across hypothalamus oligodendrocyte development.** (a) 3D UMAP plot (which is shown at a different angle and magnification than in Fig. 1 to highlight oligodendrocytes) showing distribution of oligodendrocyte-lineage cells of the hypothalamus. (b) Re-clustering of hypothalamic oligodendrocyte-lineage cells showed a clear distinction of oligodendrocyte-precursor cells (OPC), newly formed oligodendrocytes (NFO) and mature oligodendrocytes (MO); with clusters being highlighted by *Pdgfra* (OPC), *Fyn* (NFO), and *Mobp* (MO) expression. (c) Heatmap showing dynamics of gene expression across oligodendrocyte development. (d) Hypothalamus MO cluster was compared to the MO populations identified from previously published cortical scRNA-Seq and adult hypothalamus, dataset, and cross-checked against the Allen Brain Atlas (ABA) *in situ* atlas. (e) Violin plot showing *Pcsk1n* gene expression between the cortex and hypothalamus. (f) ABA *in situ* data showing *Pcsk1n* mRNA in DIG reaction (top) and colorimetric expression (bottom). (g) Violin plot showing *Cbx3* gene expression between the cortex and hypothalamus. (h) ABA *in situ* data showing raw (top) and normalized (bottom) *Cbx3* mRNA distribution. Scale bar = 1.5 mm.

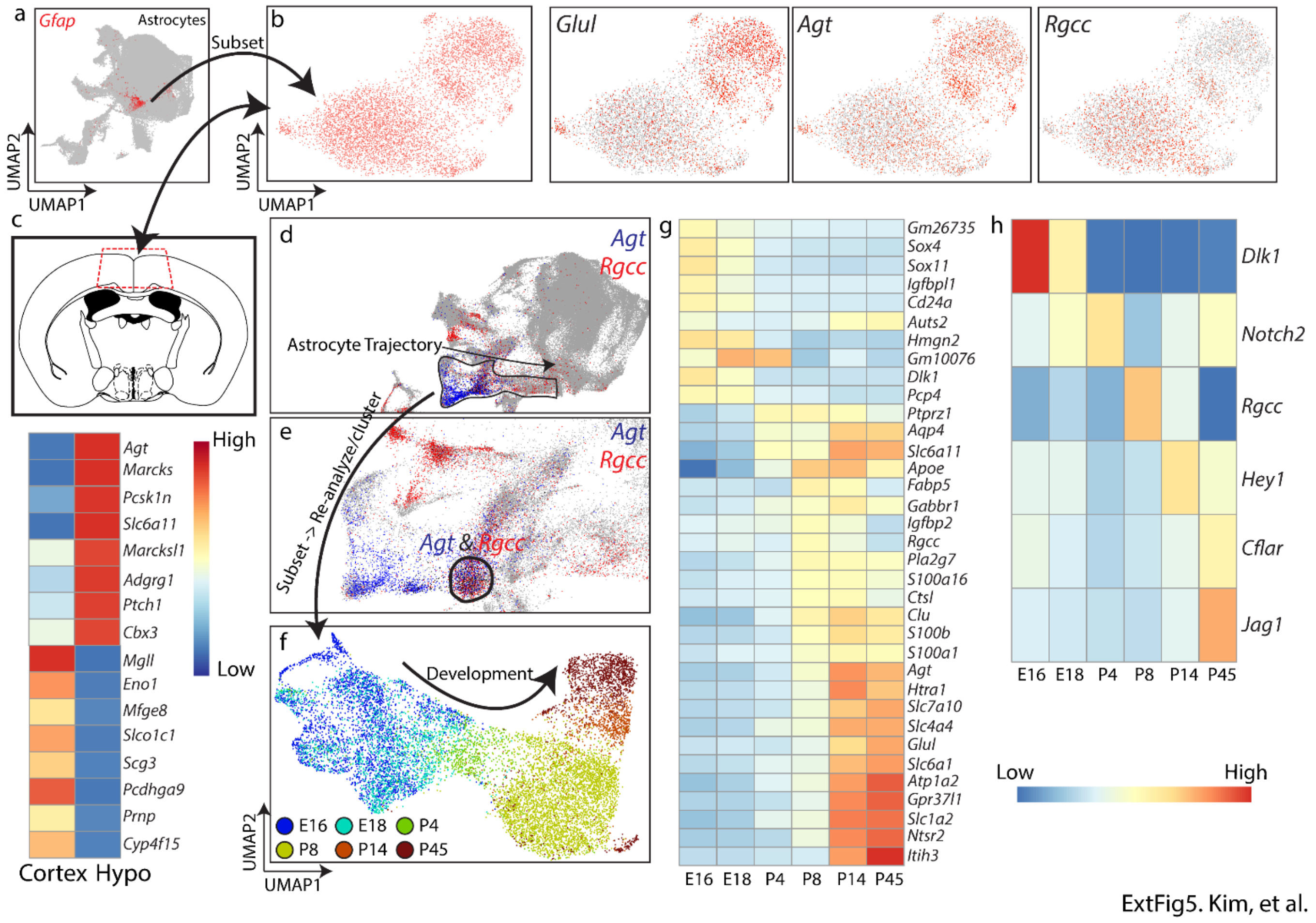


**Extended Figure 5. Dynamics of gene expression across hypothalamus astrocyte development.** (a) UMAP plot showing astrocytes highlighted with *Gfap* mRNA. (b) Astrocyte population was reclustered, and two distinct clusters were identified with varying expression levels of *Glul* and *Agt*, labelling more mature astrocyte clusters, and clusters with high *Rgcc* expression. (c) Hypothalamic astrocytes were compared to cortical astrocytes (top), with heatmap showing genes that are differentially expressed between these regions, a subset of which have been previously identified[10]. (d-f) *Agt* and *Rgcc* were plotted on the entire hypothalamic dataset (d). In the astrocyte differentiation trajectory identified in the scCoGAPS pattern highlighted by the black line, a subset of cells are identified that co-express high levels of *Agt* and *Rgcc*, representing immature astrocytes (e). (f) shows differentiation trajectory of astrocyte development from gliogenic NPCs. (g) Heatmap showing dynamics of gene expression across hypothalamic astrocyte development. (h) Heatmap showing dynamics of Notch-related signaling genes across hypothalamic astrocyte development.

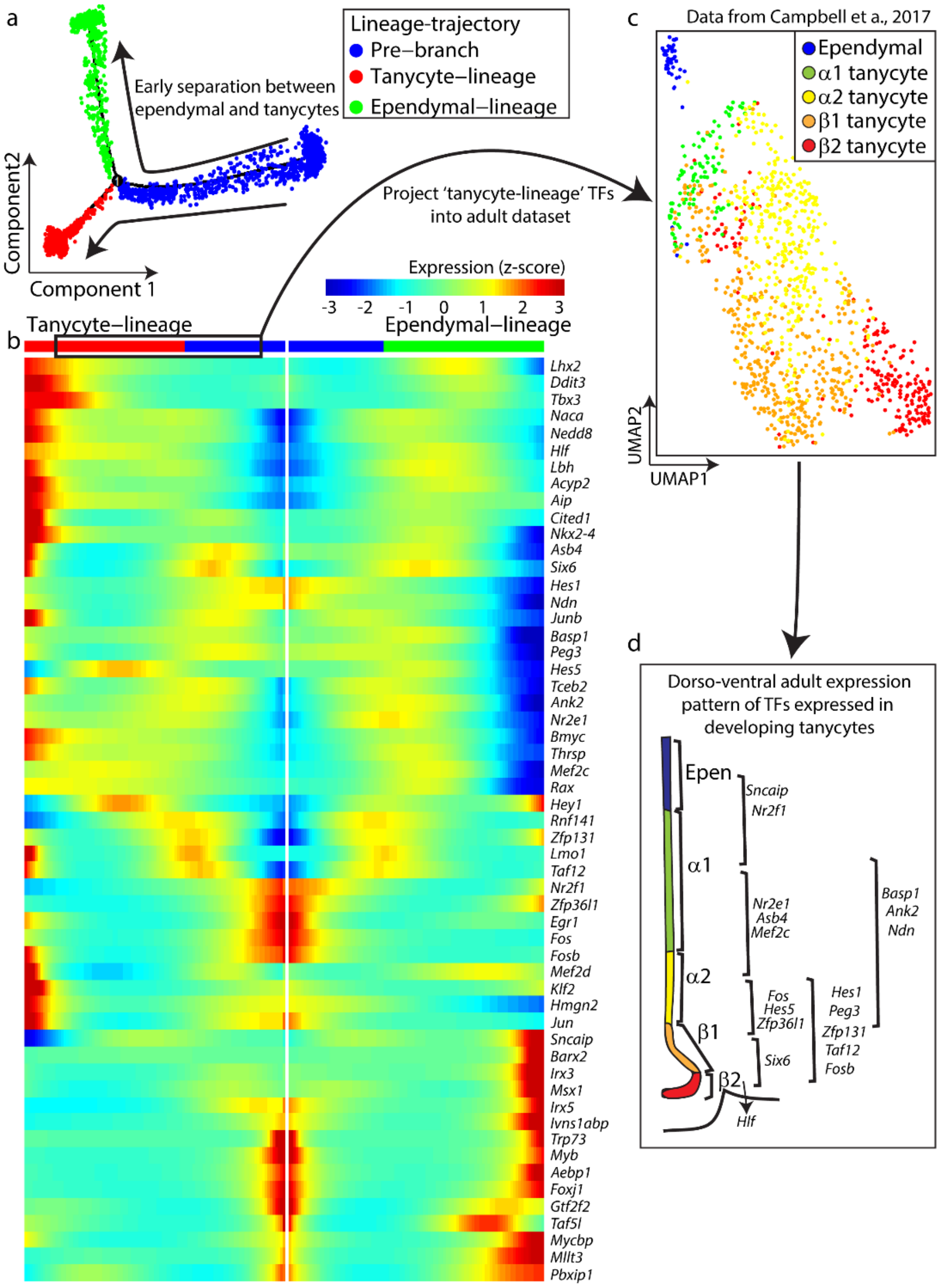


ExtFig6. Kim, et al.

**Extended Figure 6. Pseudotime analysis of differentiation of progenitors into ependymal cells and tanycytes.** (a) Pseudotime-trajectory from pre-branch (blue) to differentiation into ependymal lineage (green) or tanycyte lineage (red). (b) Pseudotime heatmap from Monocle2 BEAM analysis focusing on transcription factors that control differentiation of common progenitors into ependymal cells or tanycytes. (c) UMAP plot showing distribution of ependymal cells and subtypes of tanycytes in previously published scRNA-Seq data from adult mediobasal hypothalamus[13]. (d) Schematic showing gradients of transcription factors that are selectively expressed in different domains along the dorsoventral axis of the mediobasal hypothalamus.

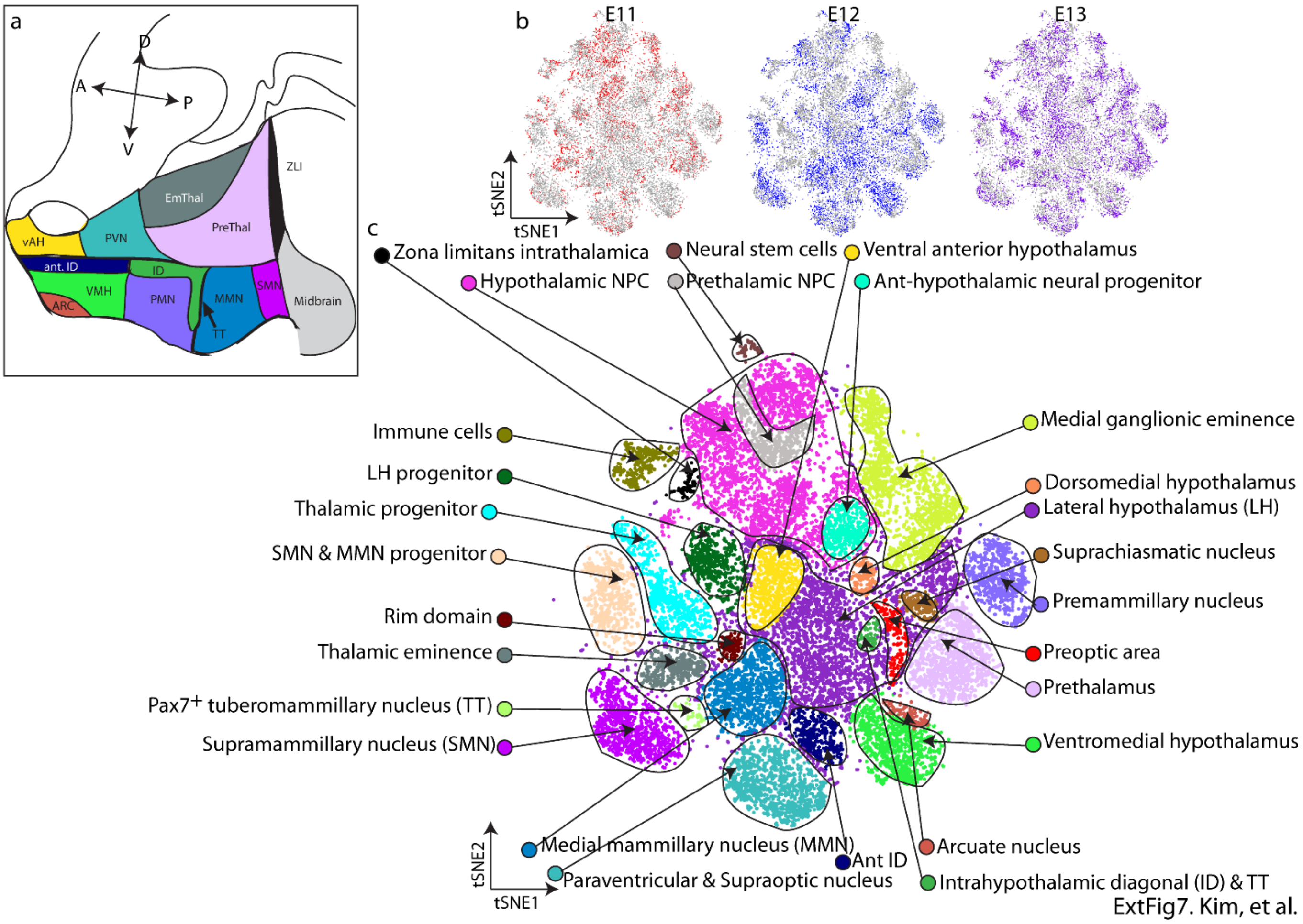


**Extended Figure 7. tSNE plot showing spatial divisions of the developing diencephalon (prethalamus and hypothalamus) and adjacent brain regions.** (a) Sagittal plane of embryonic brain highlighting the developing diencephalon. (b) tSNE plot highlighting cells from E11, E12 and E13. (c) tSNE plot of E11-E13 developing diencephalon and adjacent regions.

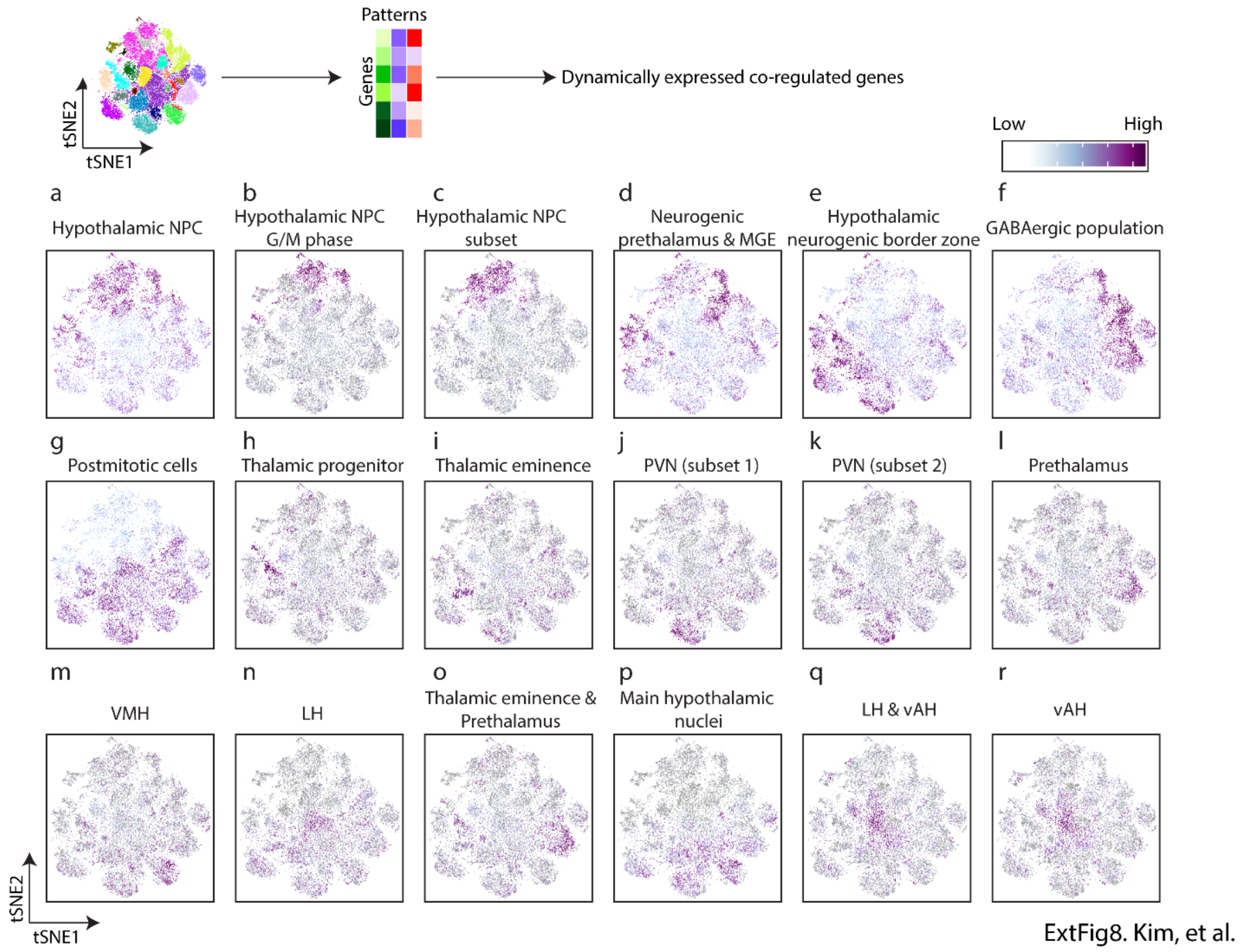


**Extended Figure 8. scCoGAPS analysis on developing hypothalamus reveals patterns capturing development or specific subregions of the hypothalamus.** scCoGAPS patterns are projected into the tSNE plot of the aggregate E11-13 dataset of the developing hypothalamus capturing hypothalamic NPC (a), G2/M phase hypothalamic NPC (b), NPCs that lack expression of hypothalamic region-specific markers (c), neurogenic prethalamic and MGE NPC (d), neurogenic border zone hypothalamic NPC (e), GABAergic NPC (f), postmitotic cells (g), thalamic NPC (h). thalamic eminence (i), PVN (subset 1)(j), PVN (subset 2)(k), prethalamus (l); VMH (m), LH (n). thalamic eminence and prethalamus (o), multiple hypothalamic nuclei (p); LH and vAH (q), vAH (r).

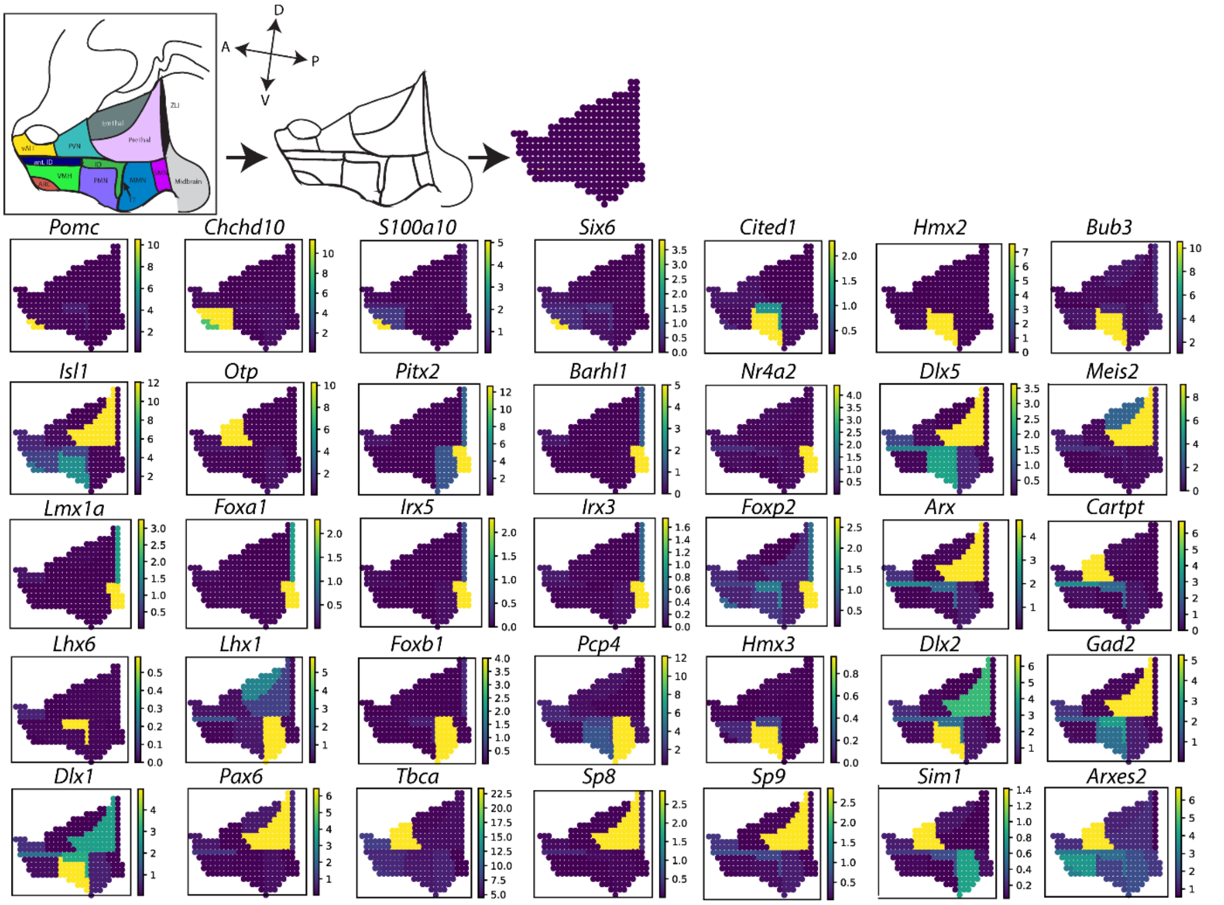


ExtFig9. Kim, et al.

**Extended Figure 9. 2D sagittal atlas of the developing diencephalon (hypothalamus and prethalamus) and projection of pattern-specific genes demarcating individual diencephalic regions.**

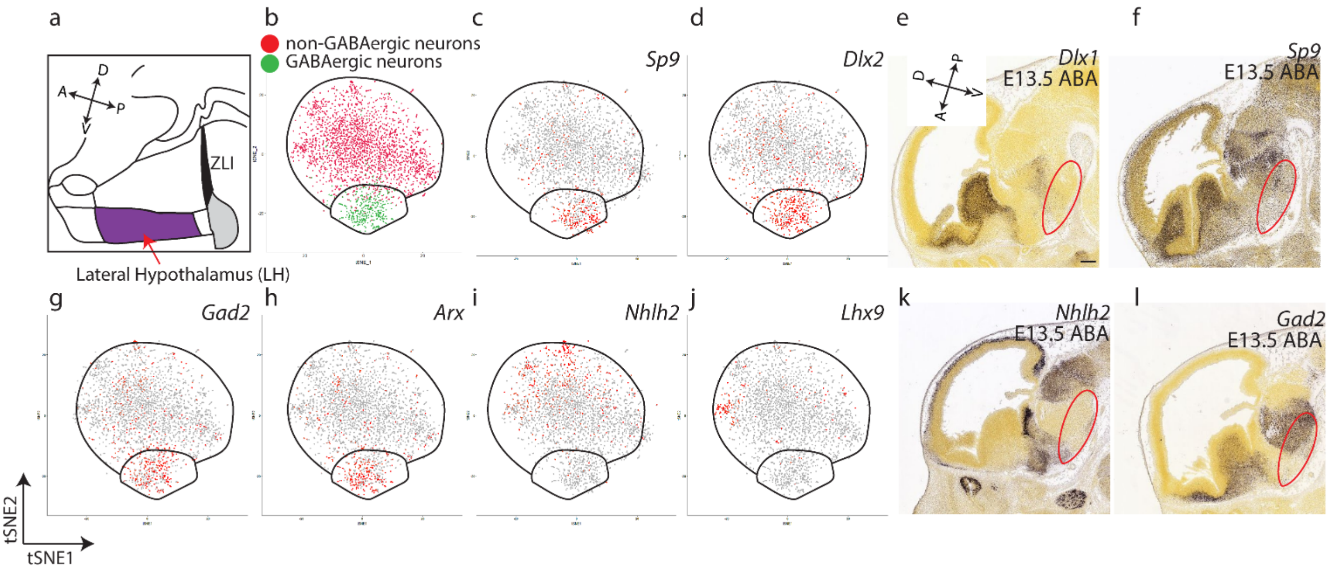


ExtFig10. Kim, et al.

**Extended Figure 10. Detailed analysis of LH shows GABAergic and non-GABAergic clusters.** (a) Parasagittal schematic highlighting the LH at E12. (b-d, g-j) tSNE plot showing clusters (b), *Sp9* (c), *Dlx2* (d), *Gad2* (g), *Arx* (h), *Nhlh2* (i), *Lhx9* (j). (e-f, k-l) *In situ* hybridization from the Allen Brain atlas showing *Dlx1* (e), *Sp9* (f), *Nhlh2* (k), *Gad2* (l). Scale bar = 0.2 mm.

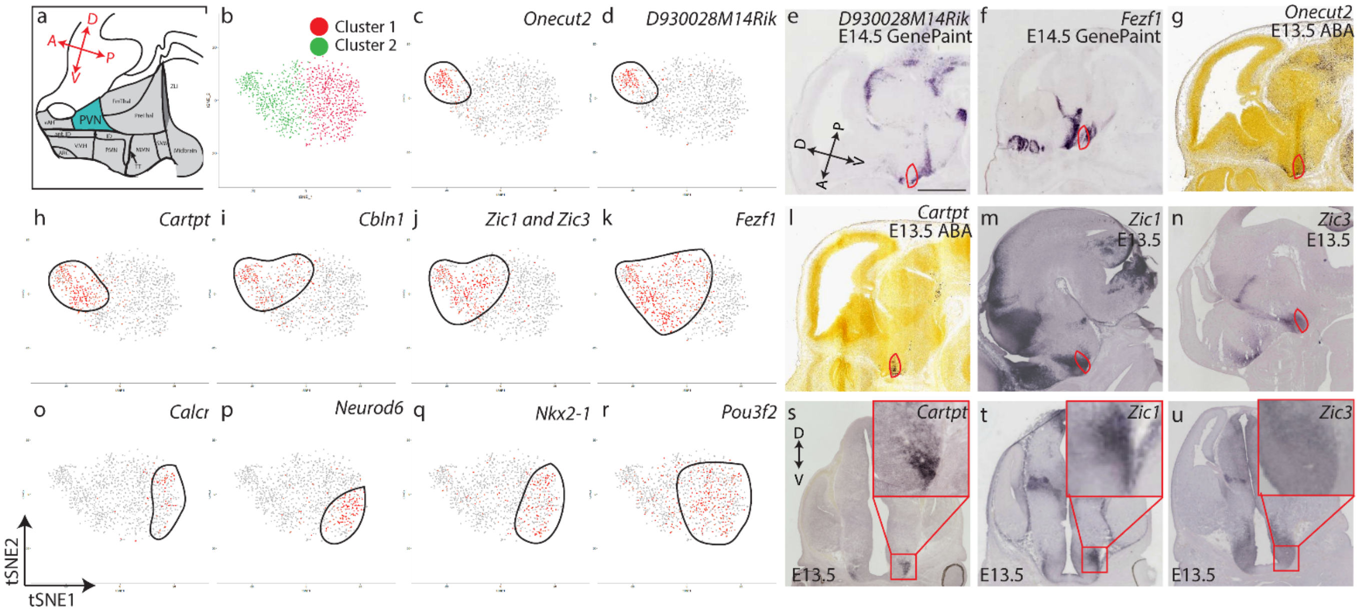


ExtFig11. Kim, et al.

**Extended Figure 11. Detailed analysis of PVN and SON during embryonic development.** (a) Schematic highlighting the PVN/SON region. (b-d, h-k, o-r) tSNE plot showing clusters (b), *Onecut2* (c), *D930028M14Rik* (d), *Cartpt* (h), *Cbln1* (i), *Zic or Zic3* (j), *Fezf1* (k), *Calcr* (o), *Neurod6* (p), *Nkx2-1* (q), *Pou3f2* (r). (e-g, l-n, s-u) *In situ* hybridization from GenePaint, ABA and our own data showing *D930028M14Rik* (e), *Fezf1* (f), *Onecut2* (g), *Cartpt* (l), *Zic1* (m), *Zic3* (n), *Cartpt* – coronal plane (s), *Zic1* – coronal plane (t), *Zic3* – coronal plane (u). Scale bar = 0.55 mm (e, g, i, m, n), 1 mm (f), 0.7 mm (s, t, u).

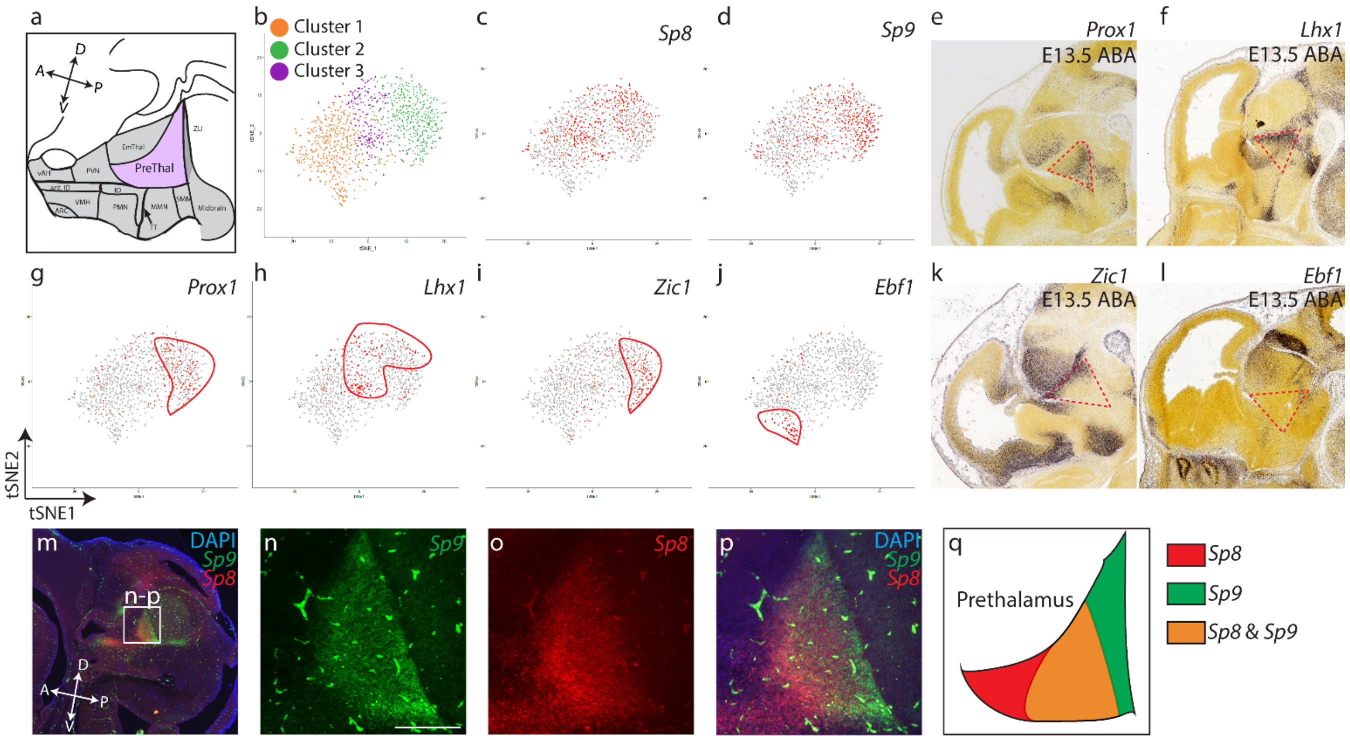


ExtFig12. Kim, et al.

**Extended Figure 12. Detailed analysis of prethalamus.** (a) Schematic highlighting the prethalamic region. (b-d, g-j) tSNE plot showing clusters (b), *Sp8* (c), *Sp9* (d), *Prox1* (g), *Lhx1* (h), *Zic1* (i), *Ebf1* (j). (e-f, k-l) *In situ* hybridization showing *Prox1* (e), *Lhx1* (f), *Zic1* (k), *Ebf1* (l). (m-q) RNAscope shows *Sp9*-positive (n) and *Sp8*-positive (o) regions of the prethalamus, and a region that co-express *Sp9* and *Sp8* (m, p, q). Scale bar = 0.2 mm.

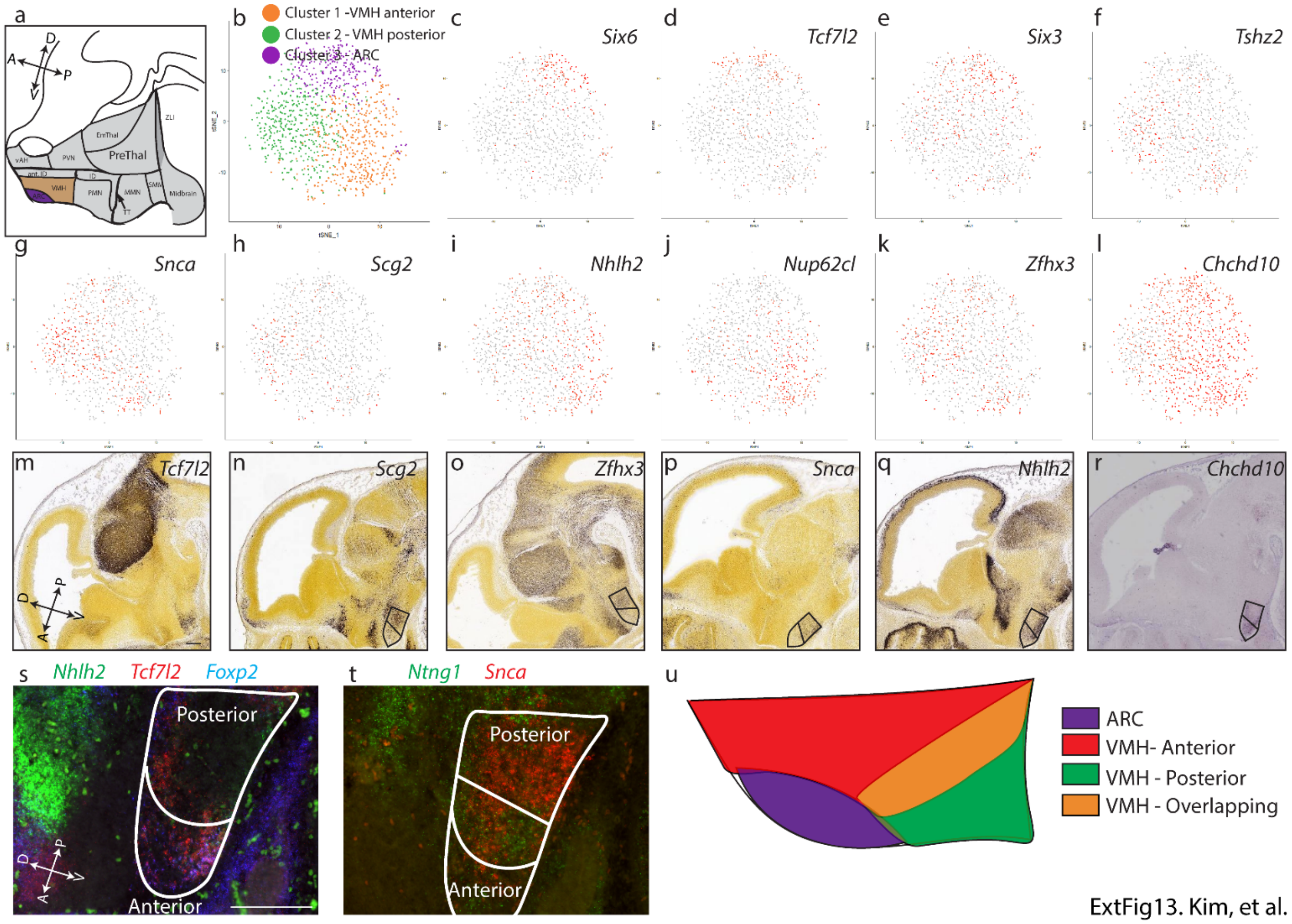


**Extended Figure 13. Detailed analysis of VMH.** (a) Schematic highlighting the VMH region. (b-l) tSNE plot showing clusters (b), *Six6* (c), *Tcf7l2* (d), *Six3* (e), *Tshz2* (f), *Snca* (g), *Scg2* (h), *Nhlh2* (i), *Nup62cl* (j), *Zfhx3* (k), *Chchd10* (l). (m-r) *In situ* hybridization from the Allen Brain atlas and GenePaint showing *Tcf7l2* (m), *Scg2* (n), *Zfhx3* (o), *Snca* (p), *Nhlh2* (q), *Chchd10* (r). RNAscope showing anterior (s) or posterior (t) regions of the VMH (white line). (u) Schematic highlighting the anterior and posterior regions of the developing VMH. Scale bar = 0.2 mm.

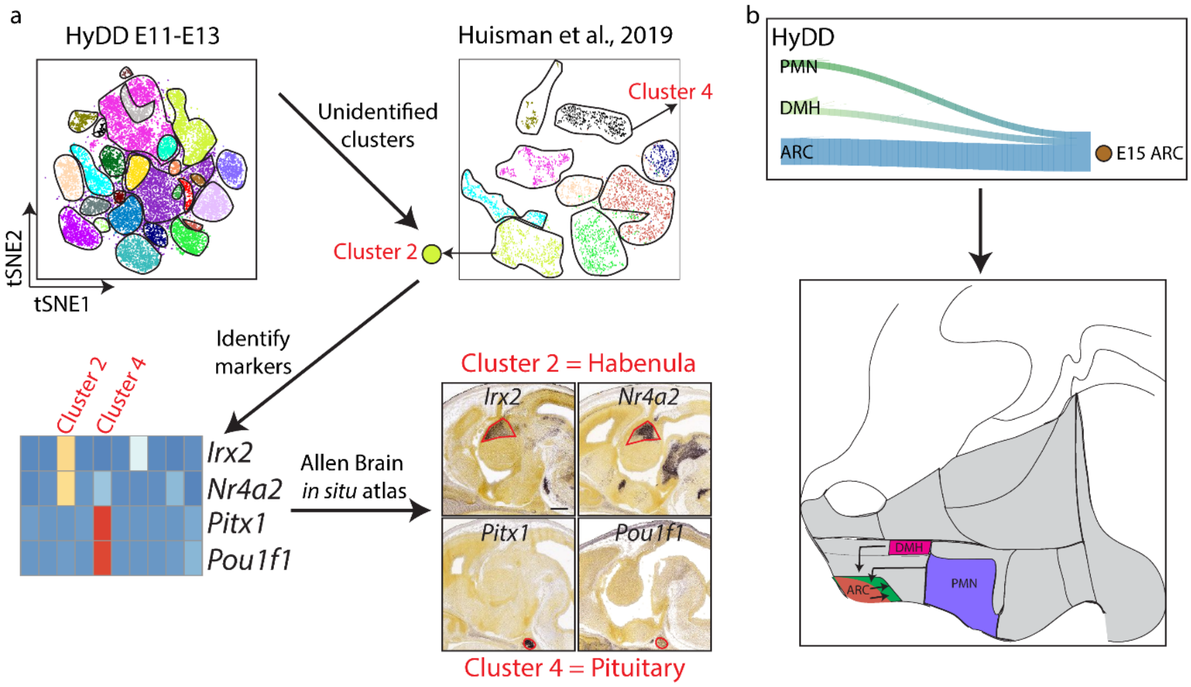


ExtFig14. Kim, et al.

**Extended Figure 14. Identification of un-matched clusters in public E15 hypothalamus scRNA-Seq dataset.** (a) Key markers of two clusters from Huisman et al., 2019[22] that were not identified by the HyDD dataset (top) were visualised in the Allen Brain *in situ* atlas (bottom). Unmatched clusters were derived from extra-hypothalamic regions of habenula and pituitary (bottom). (b) E15 arcuate nucleus showed mixed developmental origin (top), with most populations derived from embryonic arcuate nucleus, with some cells derived from from the PMN and DMH (bottom).

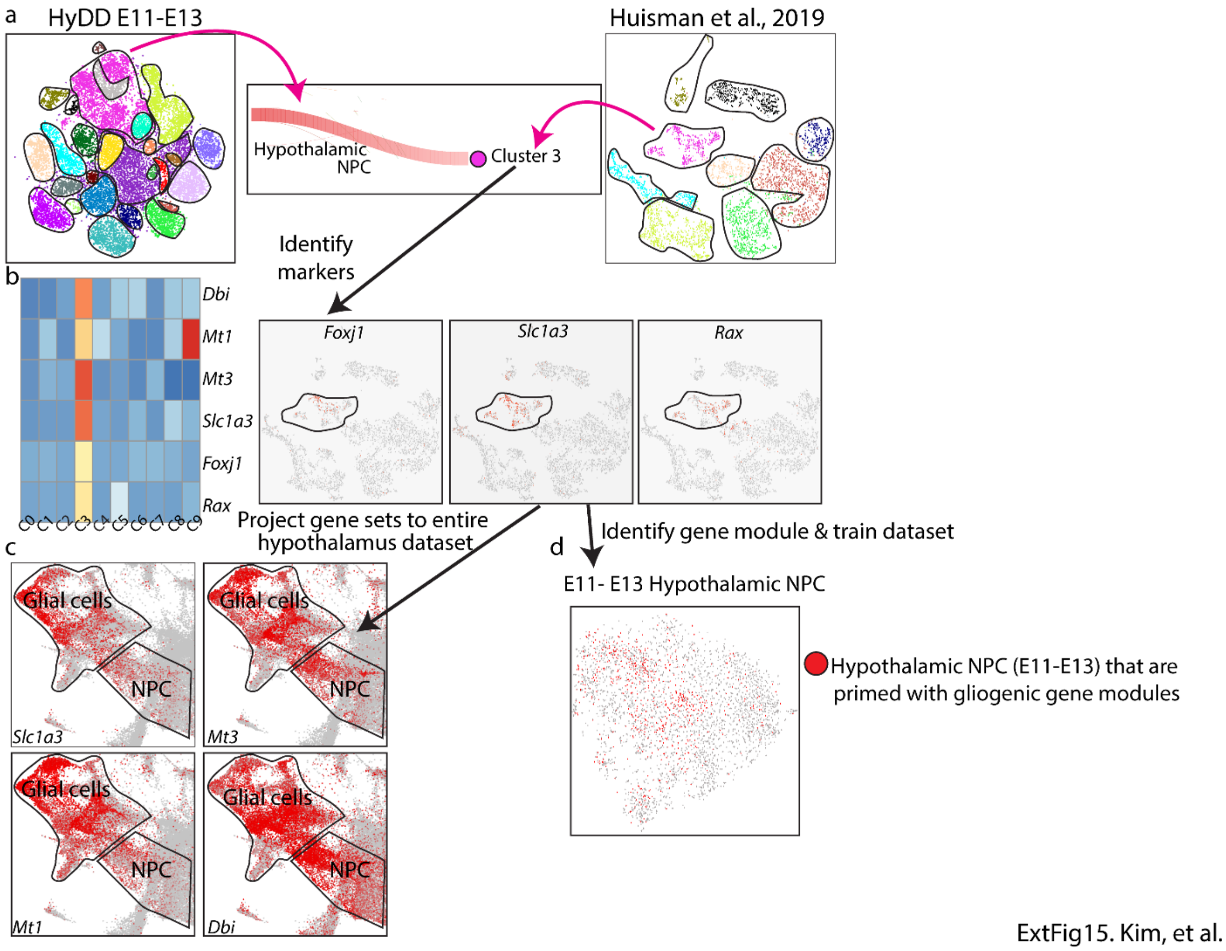


**Extended Figure 15. Identification of hypothalamic progenitors primed with gliogenic genes.** (a) Cluster 3 from Huisman et al., 2019[22] was closely related to hypothalamic NPC identified in the HyDD dataset. (b) Heatmap showing gliogenic markers of cluster 3 (left), as well as plots showing markers of ependymal cells (*Foxj1*), astrocyte/tancytes (*Slc1a3*), and tanycytes (*Rax*). (c) Visualization of key cluster 3 markers on the entire hypothalamic dataset shows strong gene expression in glial cells and sub-populations of NPC that map to glial cell differentiation trajectories. (d) Training HyDD hypothalamic NPC with identified genes of cluster 3 shows sub-populations of NPCs that selectively express gliogenic genes.

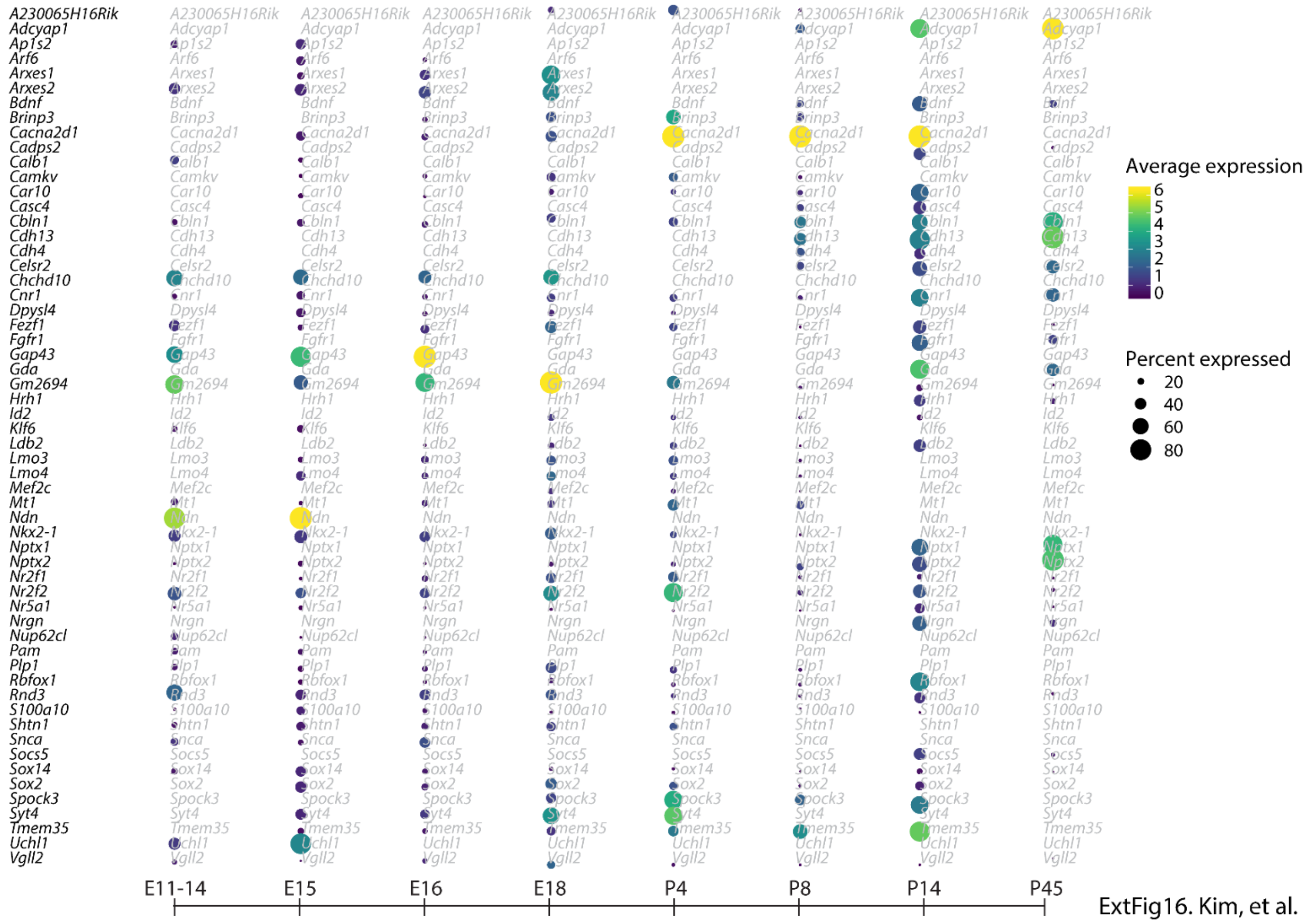


**Extended Figure 16. Using molecular stepping stones to analyze development of the VMH.** A dot plot showing key genes that can demarcate VMH across developmental time points - some genes are expressed in multiple other regions (i.e. *Nkx2-1*), others are highly specific to the VMH (i.e. *Nr5a1*), and most genes show both varying cellular levels of expression and percentages of expression in the identified VMH across development in scRNA-Seq dataset.

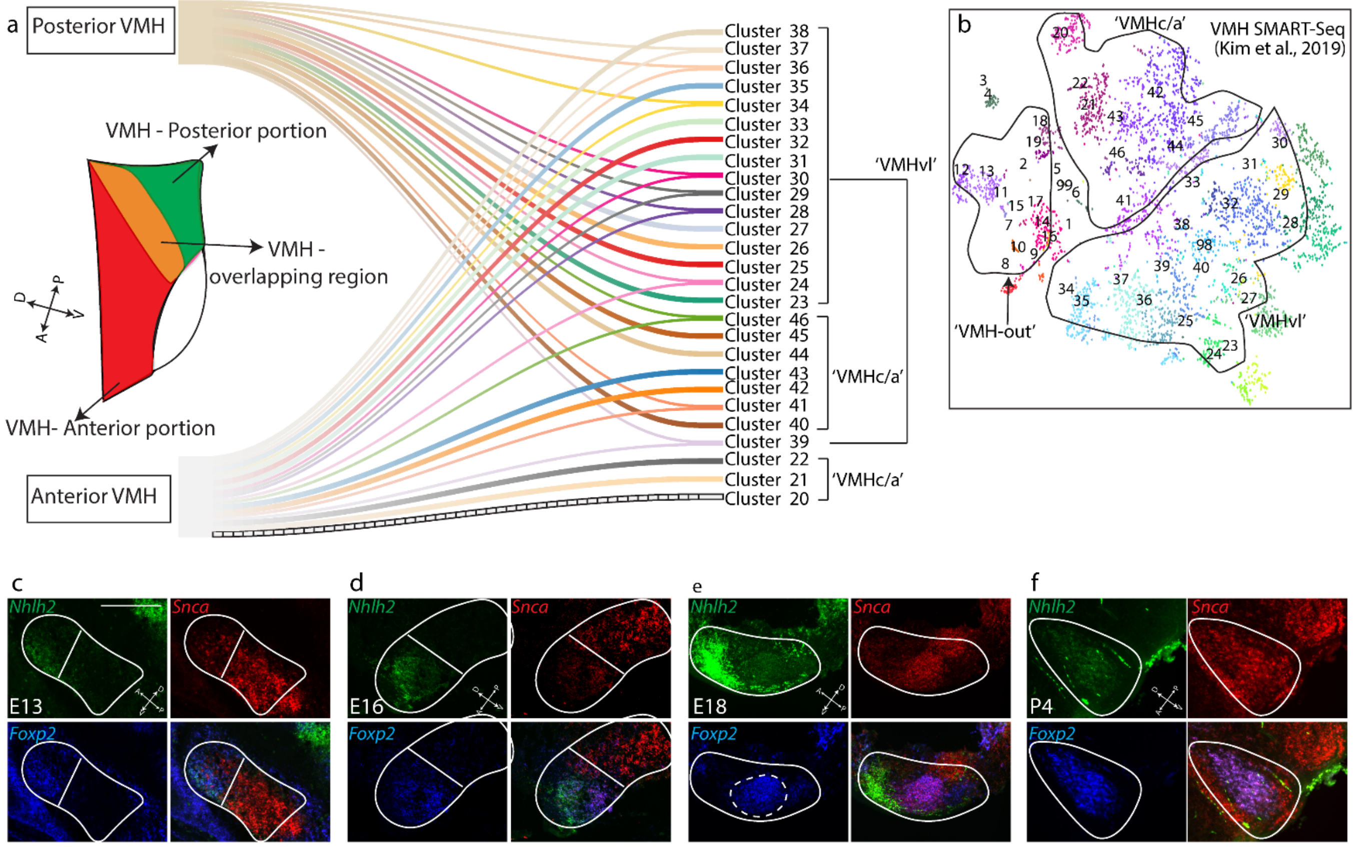


ExtFig17. Kim, et al.

**Extended Figure 17. Developmental origin of adult VMH neurons.** (a) VMH clusters obtained from Kim et al., 2019[21] were trained using markers that selectively label anterior and posterior domains of the developing VMH to identify their developmental origin. (b) tSNE plot showing the distribution of VMH SMART-Seq from Kim et al., 2019[21]. VMH clusters are derived from the original paper. (c-f) RNAscope shows distinct anterior-posterior regions of the VMH at E13 (c) and E16 (d) based on the anterior VMH markers *Nhlh2* and *Foxp2* and the posterior VMH marker *Snca*. From E18 (e) onwards, anterior VMH regions start to intermingle with posterior VMH regions. At P4 (f), anterior and posterior VMH regions are extensively intermixed. Scale bar = 0.2 mm (c), 0.25 mm (d), 0.3 mm (e) 0.4 mm (f).

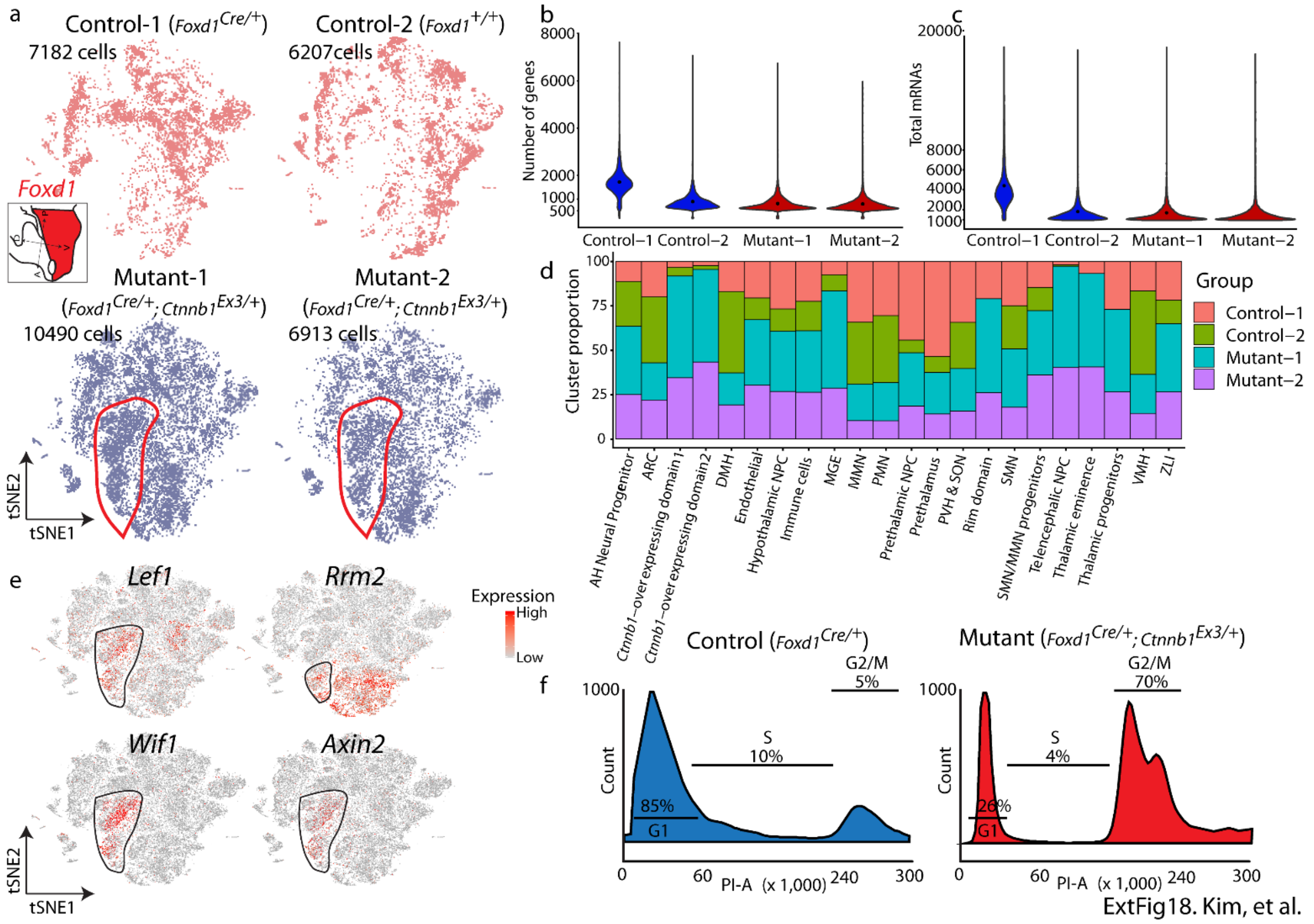


**Extended Figure 18. Overview of cell clusters in constitutively active *Ctnnb1* mutants.** (a) tSNE showing distribution of cells in individual scRNA-Seq libraries. Note clusters (red line) that only exist in mutant groups. (b) Violin plot showing distribution of mean (black dot) and of number of genes in individual scRNA-Seq libraries. (c) Violin plot showing distribution of mean (black dot) and number of total mRNAs (UMI) in individual scRNA-Seq libraries. (d) Bar graph showing the distribution of individual scRNA-Seq libraries. (e) tSNE plot showing high levels of *Lef1*, *Rrm2*, *Wif1* and *Axin2* in the mutant line. (f) Flow cytometry data in combination with propidium iodide (PI) staining to show cells at different stages of the cell cycle in control (left) and mutant samples (right).

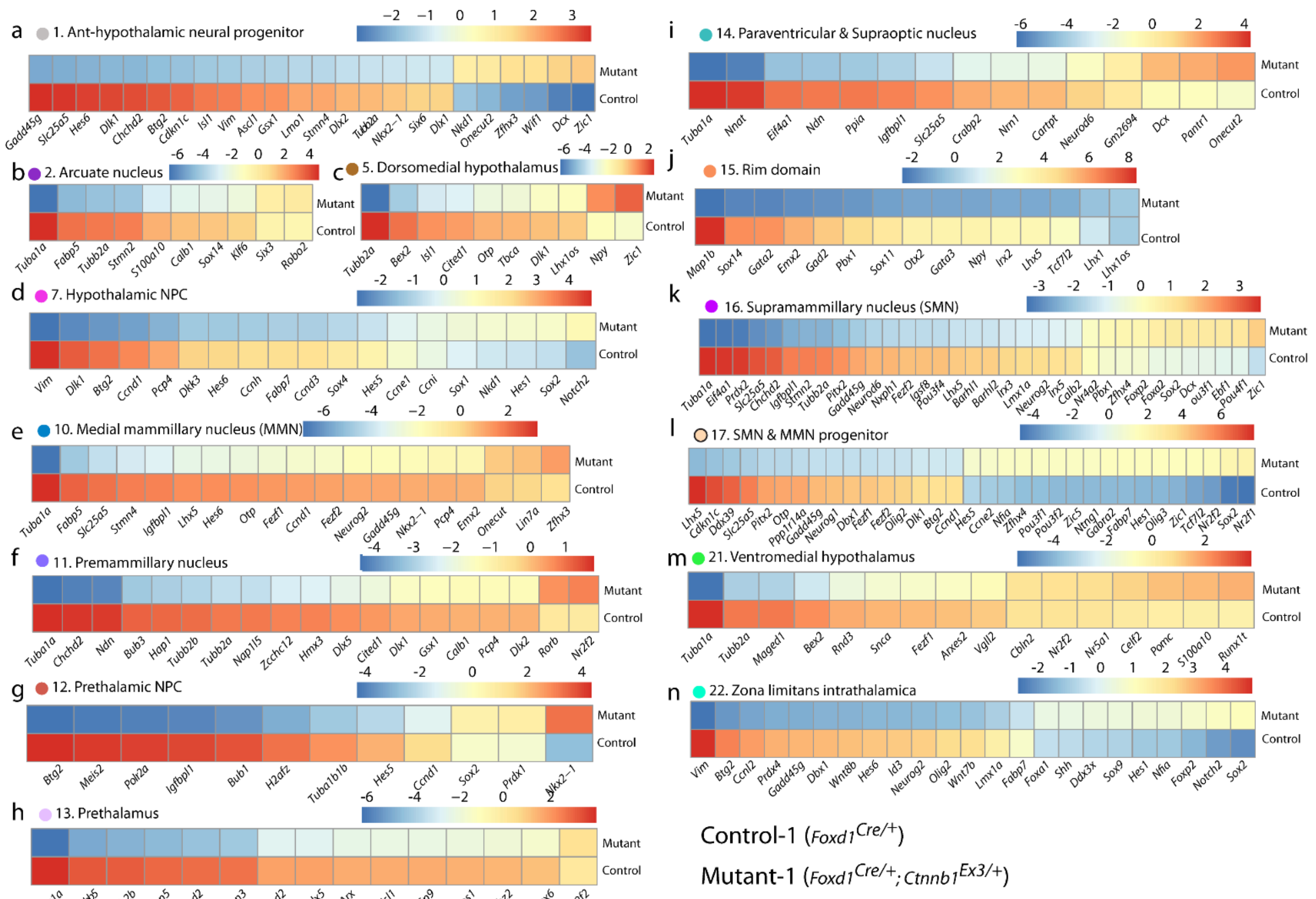


ExtFig19. Kim, et al.

**Extended Figure 19. Heatmap showing differential gene expression between control and constitutively active *Ctnnb1* mutants.** Heatmap showing top different genes between control and constitutively active *Ctnnb1* mutants in different hypothalamic regions.

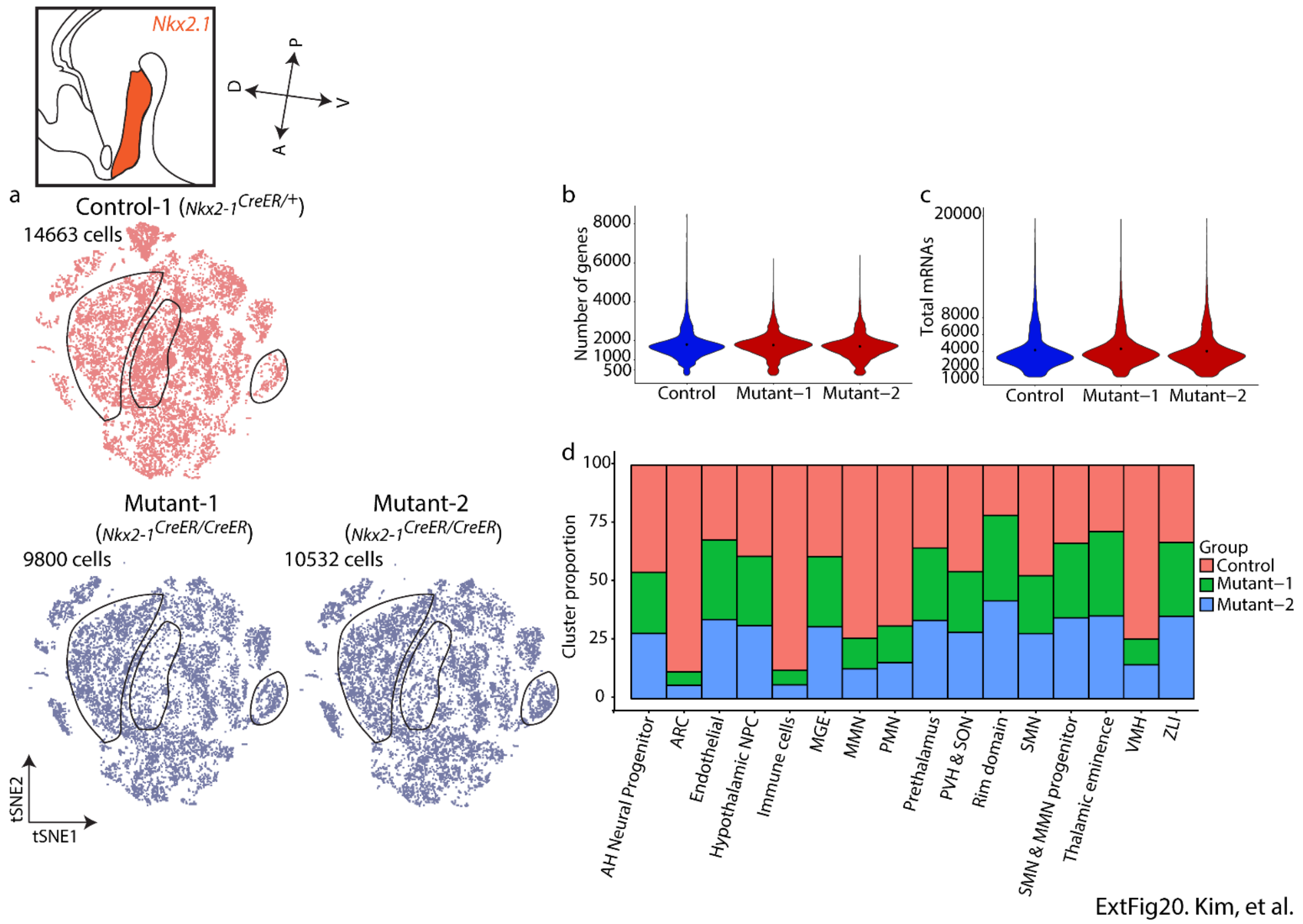


**Extended Figure 20. Overview of individual cell clusters in *Nkx2-1* mutants.** (a) tSNE showing distribution of cells in individual scRNA-Seq libraries. Note clusters (black lines) that are higher/lower in mutant groups.(b) Violin plot showing distribution of mean (black dot) and number of genes in individual scRNA-Seq libraries. (c) Violin plot showing distribution of mean (black dot) and number of total mRNAs (UMI) in individual scRNA-Seq libraries. (d) Bar graph showing distribution of individual scRNA-Seq libraries.

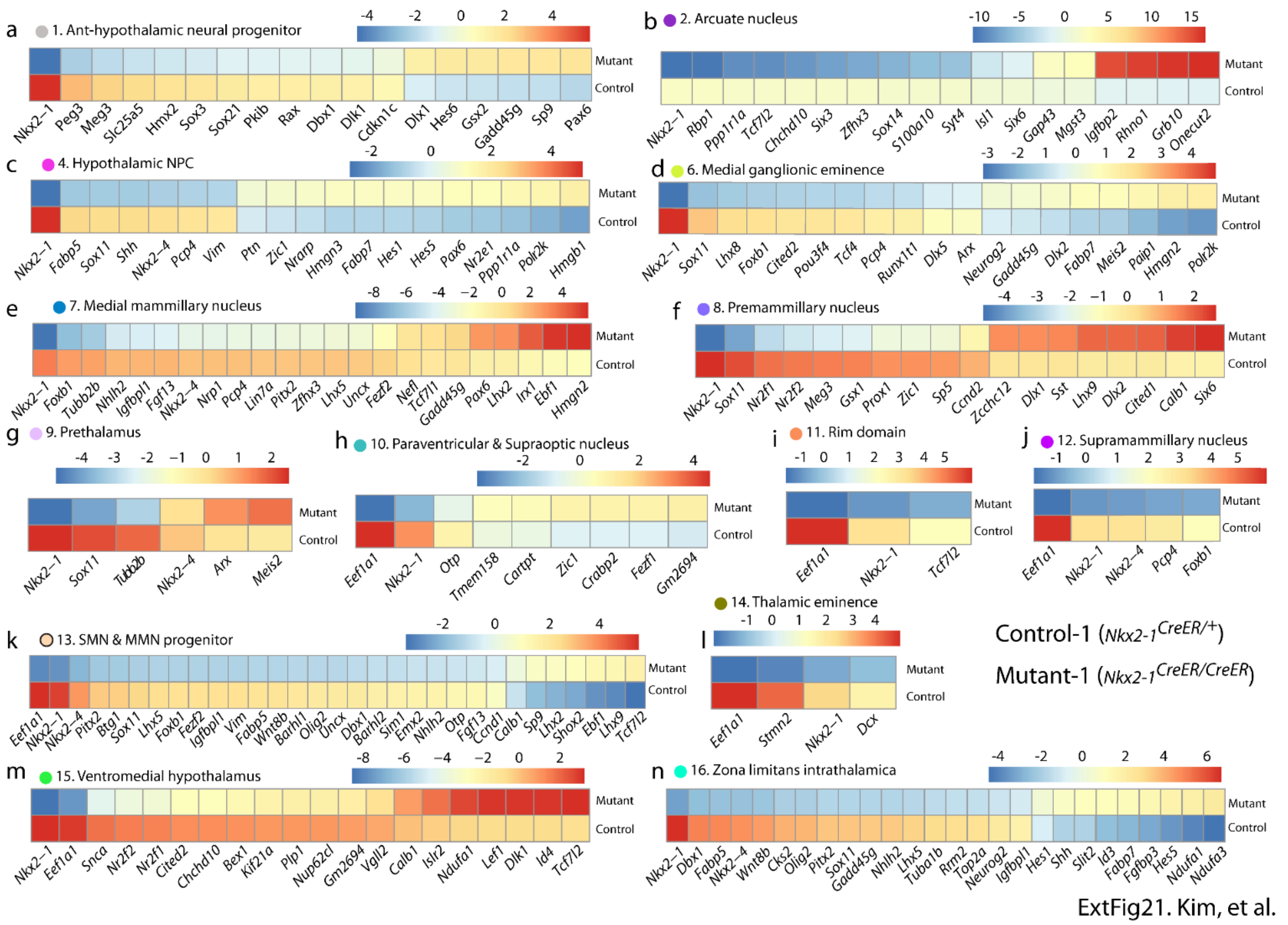


**Extended Figure 21. Heatmap showing differential gene expression between controls and *Nkx2-1* mutants.** Heatmap showing top different genes between E12.5 control and *Nkx2-1*-deficient mice in different hypothalamic regions.

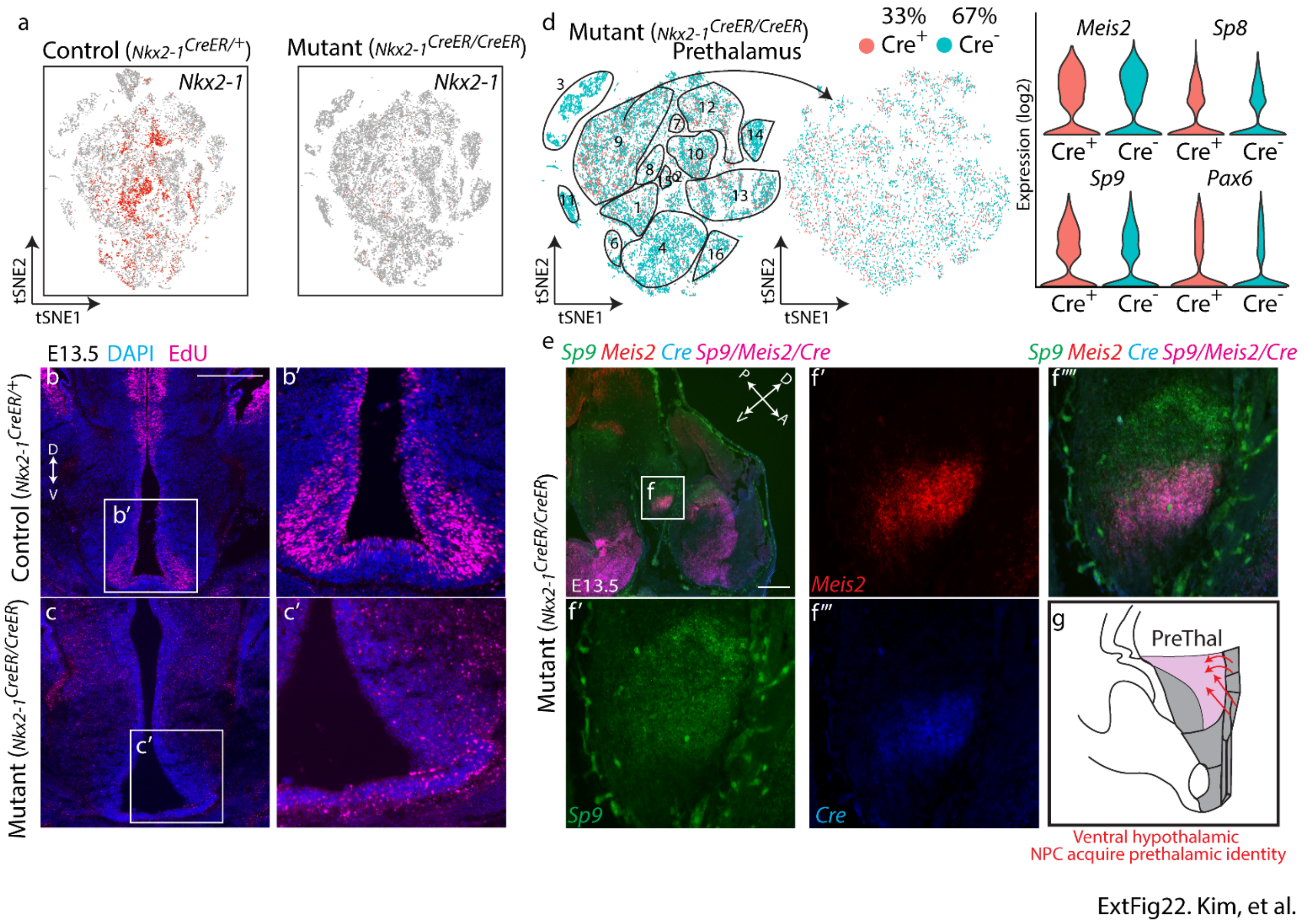


**Extended Figure 22. Characterization of *Nkx2-1* mutants.** (a) tSNE plot showing absence of *Nkx2-1* expression in mutant mice. (b-c) Images showing EdU-positive cells at E13.5 in controls (b) and mutants (c). Magnified images are shown in (b') and (c'). Note the reduction of thickness (3-fold reduction). in the ventral hypothalamus, and the increase in EdU staining in the dorsal hypothalamus. (d) tSNE plot showing cells in both controls and *Nkx2-1* mutants p expressing *Cre* (left), with *Nkx2-1*-deficient prethalamic cells highlighted (middle). Violin plots show no significant differences between expression levels of prethalamic markers between Cre-positive and Cre-negative prethalamic cells in *Nkx2-1* mutants. (e-g) RNAscope showing *Cre* expression in *Sp9*+ and *Meis2*+ prethalamic cells in *Nkx2-1* mutants (e-f), and schematic (g) showing conversion of ventral hypothalamic NPCs to prethalamic identity in the absence *Nkx2-1*. Scale bar = 0.2 mm.

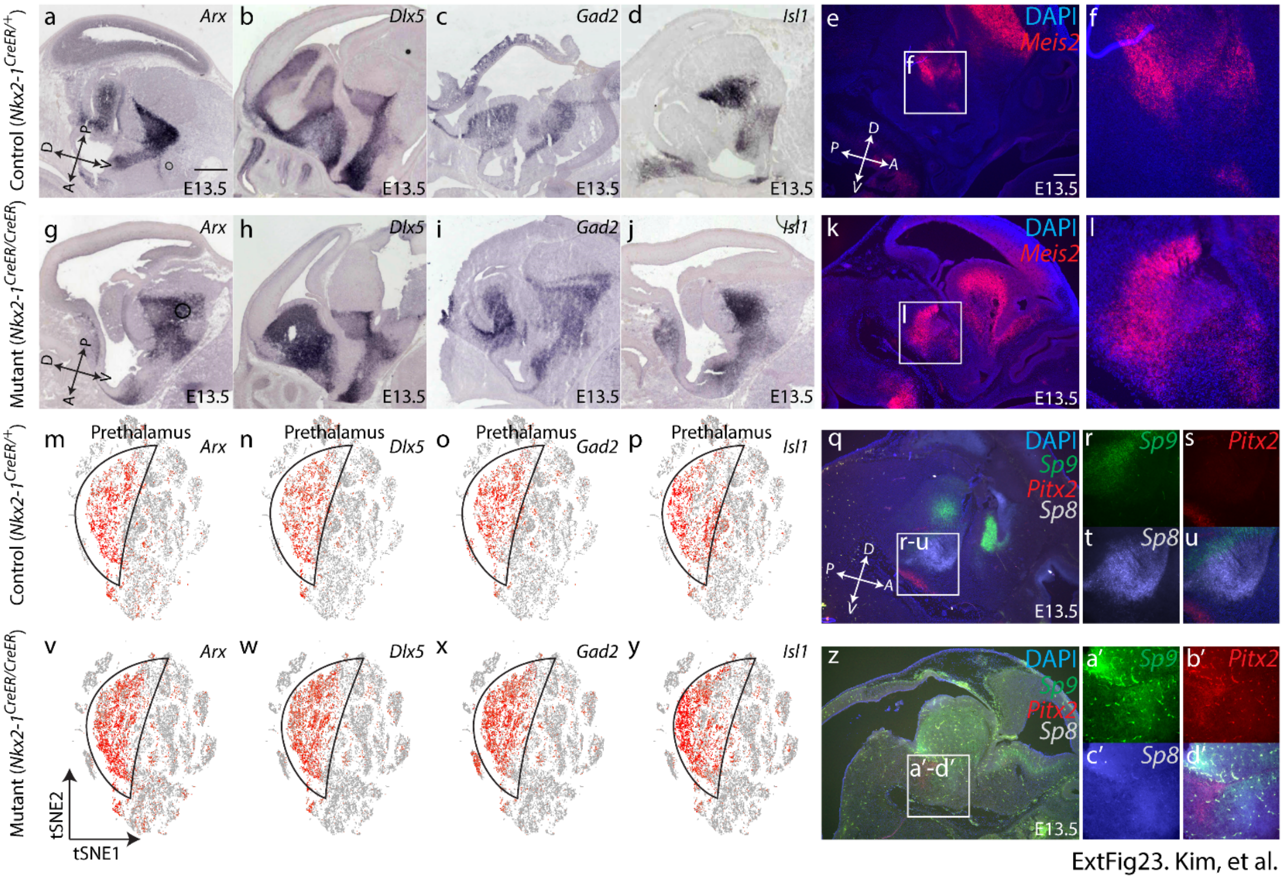

ExtFig23. Kim, et al.

**Extended Figure 23. *Nkx2-1* mutants show an increase in prethalamic cells.** (a–d, g-j) Chromogenic *in situ* hybridization showing *Arx* (a, g), *Dlx5* (b, h), *Gad2* (c, i) and *Isl1* (d, j) in control (a-d) and *Nkx2-1* mutants (g-j). (e-f, k-l) RNAScope showing *Meis2* (e-f, k-l) in control (e-f) and *Nkx2-1* mutants (k-l). (m-p, v-y) tSNE showing *Arx* (m, v), *Dlx5* (n, w), *Gad2* (o, x) and *Isl1* (p, y) in control (m-p) and *Nkx2-1* mutants (v-y). Note an increase in prethalamic cells (shown by a higher cluster of red-colored cells) and ventralization of the dorsal portion of the diencephalon. (q-u, z-c') RNAScope showing *Sp9* (r, a'), *Pitx2* (s, b') and *Sp8* (t, c') in control (q-u) and mutant (z-c'). Scale bar = 0.2 mm.

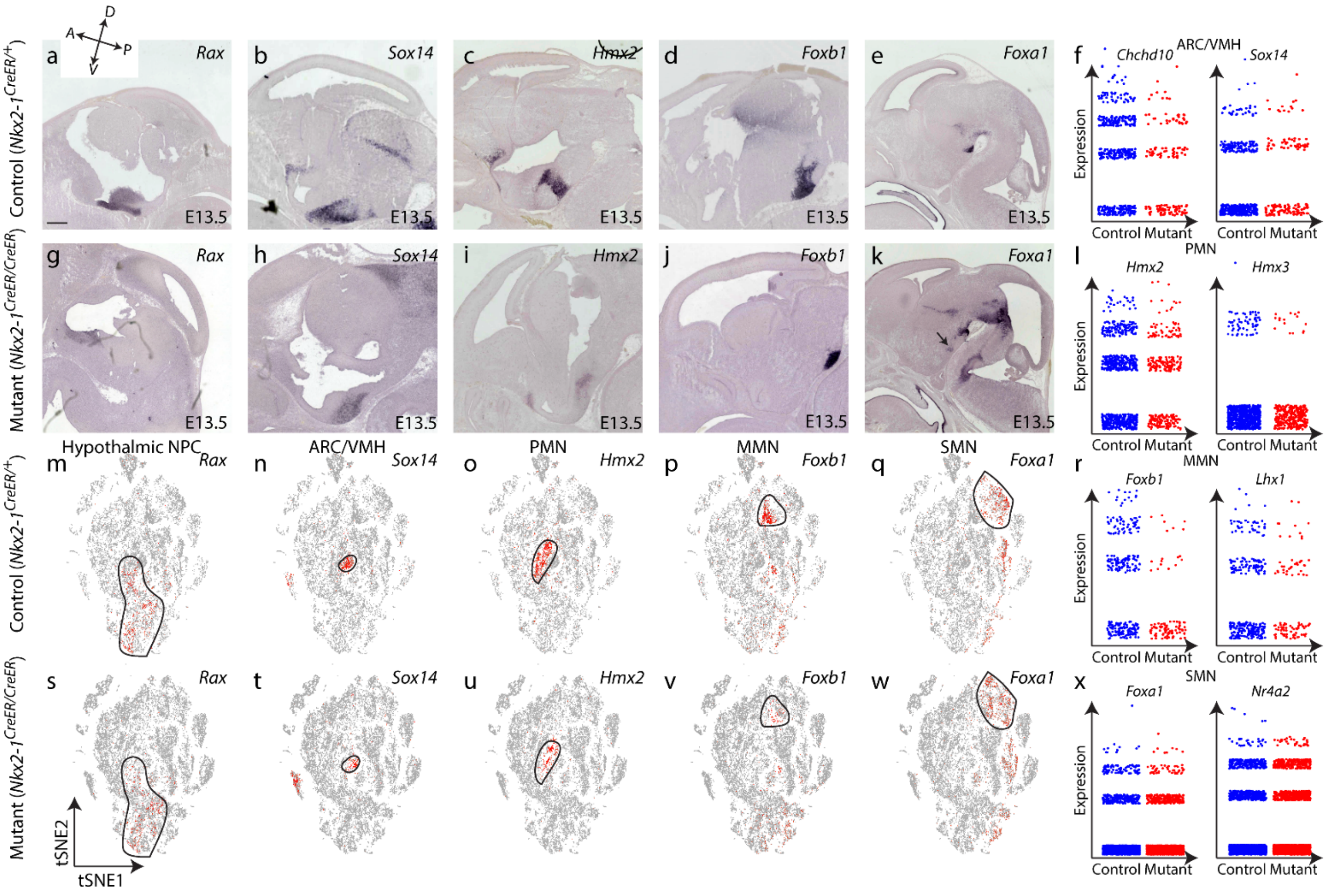


**Extended Figure 24. *Nkx2-1* mutant line shows a reduction in Nkx2.1-expressing posteroventral hypothalamic structures with the exception of the supramammillary nucleus.** (a–e, g-k) Chromogenic *in situ* hybridization showing *Rax* (a, g), *Sox14* (b, h), *Hmx2* (c, i), *Foxb1* (d, j) and *Foxa1* (e, k) in control (a-e) and *Nkx2-1* mutants (g-k). (f, l, r, x) Jitter plots of the regional marker genes (*Chchd10* and *Sox14* – ARC/VMH (f), *Hmx2* and *Hmx3* – PMN (l), *Foxb1* and *Lhx1* – MMN (r), *Foxa1* and *Nr4a2* – SMN (x)) between control and *Nkx2-1* mutants. (m–q, s-w) tSNE showing *Rax* (m, s), *Sox14* (n, t), *Hmx2* (o, u), *Foxb1* (p, v) and *Foxa1* (q, w) in control (m-q) and *Nkx2-1* mutants (s-w). Note a decrease in ventral hypothalamic markers including arcuate nucleus, ventromedial hypothalamus, premammillary nucleus and medial mammillary nucleus. Supramammillary nucleus shows no significant change, but some ectopic expression of supramammillary markers is observed in more anterior hypothalamic structures.

**Table legends:**

**Table S1. Differential gene expression between the OPC-NFO-MO in the hypothalamus.**

**Table S2. Differential gene expression in astrocytes during hypothalamic development.**

**Table S3. Pseudotime analysis of gene expression changes during tanycyte and ependymal cell development.**

**Table S4. Molecular markers for subregions of the developing hypothalamus, prethalamus , and other adjacent forebrain structures, between E11.5 and E13.5 used for generation of HyDD.**

**Table S5. scCoGAPS patterns (Fig. S11) and pattern weights for all 19,622 expressed genes.**

**Table S6. Differential gene expression between subregions of the LH, PVH, prethalamus and VMH between E11.5 and E13.5**

**Table S7. Differential gene expression between E12.5 control (*Foxd1*$^{Cre/+}$) and constitutively active *Ctnnb1* overexpressing mutant (*Foxd1*$^{Cre/+}$;*Ctnnb1*$^{Ex3/+}$) samples.**

**Table S8. Differential gene expression between E12.5 control (*Nkx2-1*$^{CreER/+}$) and *Nkx2-1*-deficient mutant (*Nkx2-1*$^{CreER/CreER}$) samples.**

**Table S9. Number of cells found in individual clusters of control and mutant samples.**
